# Automated design and optimization of multitarget schizophrenia drug candidates by deep learning

**DOI:** 10.1101/2020.03.19.999615

**Authors:** Xiaoqin Tan, Xiangrui Jiang, Yang He, Feisheng Zhong, Xutong Li, Zhaoping Xiong, Zhaojun Li, Xiaohong Liu, Chen Cui, Qingjie Zhao, Yuanchao Xie, Feipu Yang, Chunhui Wu, Jingshan Shen, Mingyue Zheng, Zhen Wang, Hualiang Jiang

**Author notes:** Corresponding authors. E-mail addresses (Mingyue Zheng), (Zhen Wang), (Hualiang Jiang). These authors contributed equally to this work (Xiaoqin Tan, Xiangrui Jiang, Yang He). Abbreviations*: GPCRs, G protein-coupled receptors; RNN, deep recurrent neural network; MTDNN, multitask deep neural network; D_2_R, D_2_ receptors; 5-HT_2A_R, 5-HT_2A_ receptors; 5-HT_1A_R, 5-HT_1A_ receptors; EPS, Parkinson-like extrapyramidal symptoms; TD, tardive dyskinesia; HTS, high-throughput screening; AI, artificial intelligence; QSAR, quantitative structure-activity relationship; AEs, autoencoders; GANs, generative adversarial networks; RL, reinforcement learning; MW, molecular weight; logP, Wildman-Crippen partition coefficient; TPSA, total polar surface area; DNNs, deep neural networks; LSTM, long short-term memory; t-SNE, t-distributed stochastic neighbor embedding; ECFP4, extended connectivity fingerprint 4; SA, synthetic accessibility; QED, quantitative estimate of drug-likeness; R^2^, correlation coefficient; MAE, mean absolute error; RO5, Lipinski’s rule of five; CNS, central nervous system; PCP, phencyclidine; NMDAR, N-methyl-D-aspartic acid receptor; SAR, structural-activity relationships; BPTT, backpropagation through time; ReLU, rectified linear unit; MSE, mean squared error.

## Abstract

Complex neuropsychiatric diseases such as schizophrenia require drugs that can target multiple G protein-coupled receptors (GPCRs) to modulate complex neuropsychiatric functions. Here, we report an automated system comprising a deep recurrent neural network (RNN) and a multitask deep neural network (MTDNN) to design and optimize multitargeted antipsychotic drugs. The system successfully generates novel molecule structures with desired multiple target activities, among which high-ranking compound **3** was synthesized, and demonstrated potent activities against dopamine D_2_, serotonin 5-HT_1A_ and 5-HT_2A_ receptors. Hit expansion based on the MTDNN was performed, 6 analogs of compound **3** were evaluated experimentally, among which compound **8** not only exhibited specific polypharmacology profiles but also showed antipsychotic effect in animal models with low potential for sedation and catalepsy, highlighting their suitability for further preclinical studies. The approach can be an efficient tool for designing lead compounds with multitarget profiles to achieve the desired efficacy in the treatment of complex neuropsychiatric diseases.

**Graphical abstract:** 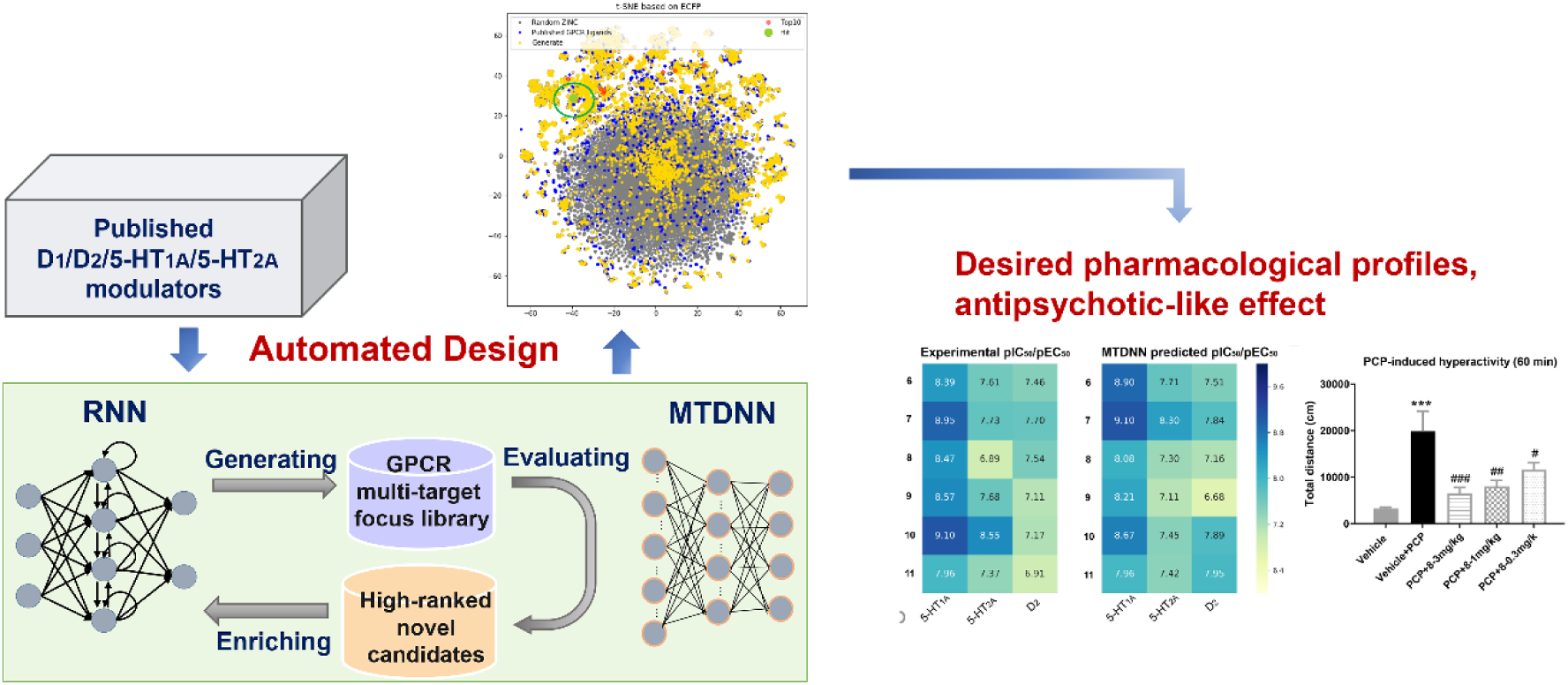

## 1. Introduction

The safety and clinical efficacy of a drug depend on its activity profile across many proteins in the proteome. Schizophrenia is a complex psychiatric disease affecting approximately 1% of the world’s population^1^, and it requires drugs targeting multiple GPCRs to modulate therapeutically complex neuropsychiatric functions^2^. The core symptoms of schizophrenia are classified into positive symptoms (delusions, hallucinations and thought disorder), negative symptoms (alogia, anhedonia and apathy) and cognitive impairment^3^. Typical antipsychotics, which mainly antagonize dopamine D_2_ receptors (D_2_R), are effective merely in treating the positive symptoms and may cause various side effects (e.g., Parkinson-like extrapyramidal symptoms (EPS), tardive dyskinesia (TD), or even fatal cardiovascular events)^4^. Atypical antipsychotics, which is characterized by low affinity for D_2_R in addition to comparatively high affinity for the serotonin 5-HT_2A_ receptors (5-HT_2A_R), reduce the EPS and TD side effects^5^. But atypical antipsychotics show unsatisfactory effects on cognitive dysfunction and the negative symptoms^6-8^ and also cause side effects, such as dyslipidemia, weight gain, diabetes and QT interval prolongation^9-11^. Recent preclinical and clinical studies have suggested that 5- HT_1A_R agonists improve the cognitive and negative symptoms of schizophrenia and reduce the EPS side effects caused by antipsychotics^12-13^. Several atypical antipsychotics such as ziprasidone, lurasidone, aripiprazole^14^, brexpiprazole^15^ and cariprazine^16^ exhibit affinity toward the 5-HT_1A_ receptor (5-HT_1A_R), but their efficacy and side effect profiles are notably different, partly owing to their differences in 5-HT_1A_R/D_2_R affinity balance. The optimal 5-HT_1A_R/D_2_R affinity balance need to be investigated to improve the effects on cognitive dysfunction and reduce the negative symptoms, the EPS and metabolic side effects. Currently, the affinities for 5-HT_1A_R of launched atypical antipsychotics is generally much lower than that for D_2_R, so it is of interest to design and evaluate compounds with higher affinity to 5-HT_1A_R than D_2_R, and with 5-HT_2A_R antagonism that may reduce the risk for EPS by increasing dopaminergic transmission. Therefore, despite the available antipsychotic drug candidates that act on dopamine and serotonin receptors, there has been an unmet demand in developing novel chemotypes with potent and balanced activities, for modulating the positive symptoms, negative symptoms and cognitive impairment with minimal side effects.

Most of the multitarget antipsychotics were discovered by serendipity but not by the rational drug design. Unfortunately, designing novel chemical compounds of multitarget profile with a specific balance is a complex and extraordinarily challenging task for medicinal chemists. The number of chemicals that are considered to be qualified drug-like molecules is estimated to be between 10^30^ and 10^60^, which is far too large even for biological high-throughput screening (HTS) assays^17^. For functional studies relying on culture-based approaches, biological activity evaluation is also impractical as it is a time-consuming and labor-intensive process^18^. Moreover, the lack of sufficient structural information regarding GPCRs has been a major obstacle in a multitude of virtual screening and rational drug design studies^19-20^. To date, several de novo drug design methods have been proposed^21-24^, most of which are based on single-molecule targets and the proposed molecules are seldom experimentally synthesized and tested.

In recent years, new approaches based on artificial intelligence (AI), especially deep learning technologies, have made an impact on the field of de novo drug design and multitarget activity prediction^25-28^. In 2012, Merck organized a Kaggle competition to find the most advanced methods for QSAR studies. Using MTDNN, the winning team achieved an accuracy 15% better than Merck’s internal baseline. It has also been shown that MTDNN models trained on multiple QSAR tasks simultaneously can outperform deep neural networks (DNNs) trained on many individual tasks separately^29^. Segler et al. used Recurrent neural networks (RNNs) to learn the syntax of the SMILES notation and established target-focused libraries by fine-tuning a pretrained network with a smaller set of compounds. This transfer learning method has been successfully used in generating a high proportion of predicted actives^30^. Popova et al. combined an RNN with reinforcement learning (RL) for de novo molecule generation and to design molecular libraries with a bias toward maximal, minimal, or a specific range of physical and chemical properties^27^. More recently, Zhavoronkov et al. have developed a deep generative model to discover potent inhibitors of discoidin domain receptors 1 (DDR1). Using this method, six potent inhibitors for DDR1 were designed and tested experimentally in 46 days, which demonstrates the potential of deep learning methods to afford rapid and effective molecular design^28^.

In this study, we developed an automated system for designing novel antipsychotic drugs based on RNN and MTDNN models (Figure 1A), which consists of three tasks: (a) generating a virtual focused library for the D_2_/5-HT_1A_/5-HT_2A_ receptors based on an RNN; (b) virtual screening with activity scoring models established by an MTDNN, owing to the significant homology (Figure S1) and a large number of ligands shared by dopaminergic and serotoninergic receptors; (c) iterative virtual optimization of the focused library by incorporating high-scoring novel compounds and eliminating the lower-scoring compounds. In each epoch, the generated molecules are evaluated and screened by various criteria, including predicted activity, synthetic accessibility, drug-likeness and structural novelty. High-ranking molecules with good drug-likeness and synthetic accessibility are sampled to expand the training set for the next iteration. Additionally, the method can be considered as a problem of reinforcement learning, where the generative model can be cast as a policy and the MTDNN model provides reward signals that can automatically generate more attractive molecules.

**Figure 1.**
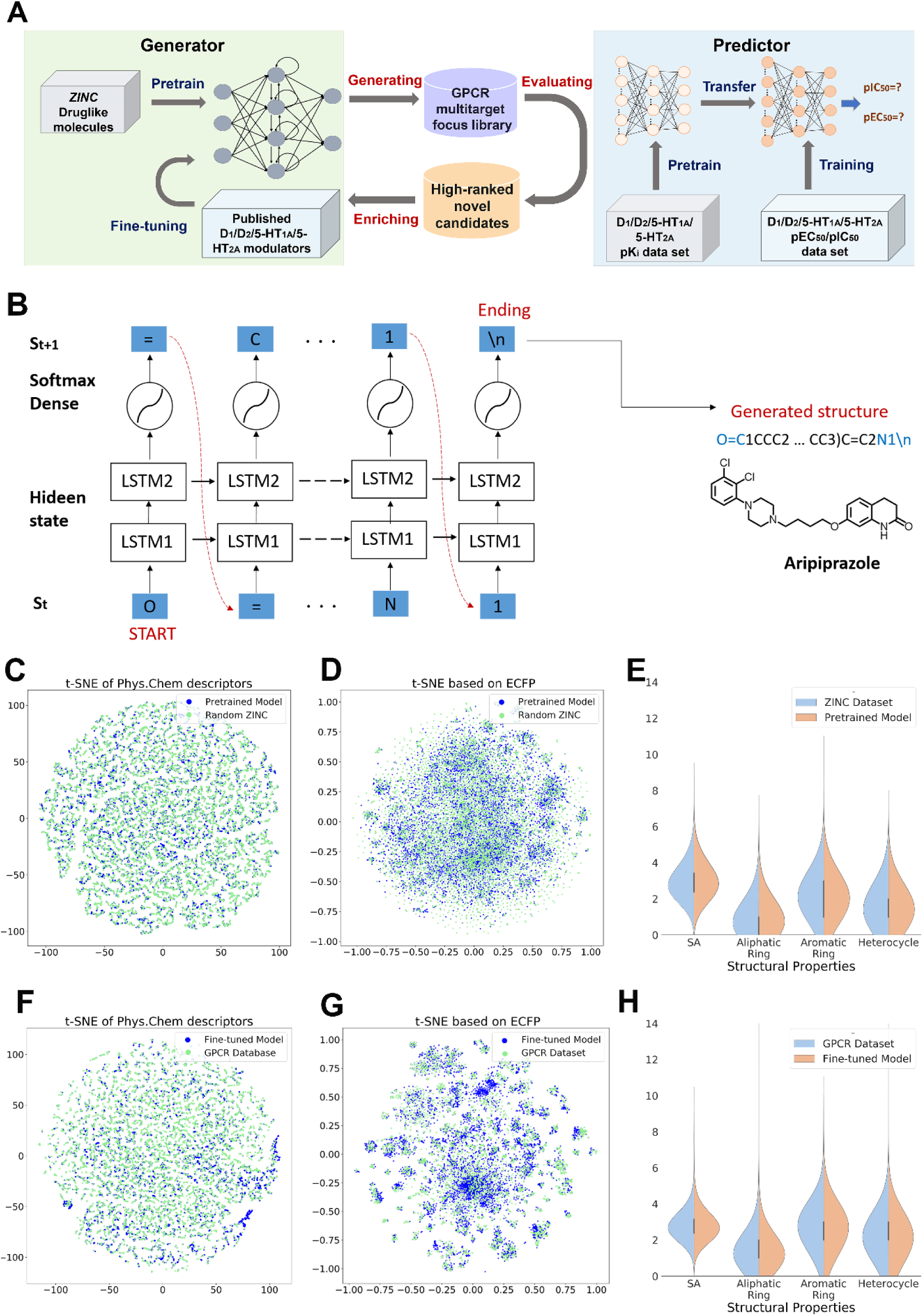
Automated molecular structure design system and distributions of the physicochemical properties of the generate molecules with the pretrained and fine-tuned model. (A) Overall workflow of de novo antipsychotic drug design cycle. (B) Model of the RNN that produces the SMILES string for aripiprazole, token-by-token. The model predicts the next token for each input token in the sequence, the ending token is “\n”. (C), (D) The physicochemical space of the molecules generated by the model pretrained with the *ZINC* data set. e Comparison of the properties including ring systems, SA scores of the molecules generated by the model pretrained with the *ZINC* data set. (F), (G) The physicochemical space of the molecules generated by the model fine-tuned with the published GPCR data set. (H) Comparison of the properties including ring systems, SA scores of the molecules generated by the model fine-tuned with the published GPCR data set.

## 2. Results and discussion

### 2.1. Deep generative model for molecular structure generation by the RNN

The computational approach for generating target-specific libraries consists of two basic steps: (a) learning SMILES by an RNN model trained with a large set of drug-like compounds, and (b) fine-tuning the RNN model based on a small library of compounds active against D_1_/D_2_/5-HT_1A_/5-HT_2A_ receptors.

Structurally, our model consists of two stacked long short-term memory (LSTM) layers, each with 1024 dimensions, regularized with a dropout layer with a dropout ratio of 0.2 (as shown in Figure 1B). To predict the next character in the SMILES strings representing molecular structures, these two layers are followed by a dense output layer and a neuron unit with a *softmax* activation function. Here, the generation of aripiprazole is presented as an example. Aripiprazole is primarily used for the treatment of schizophrenia or bipolar disorder,^14^ and it can be described by the SMILES string: “O=C1CCC2=CC=C (OCCCCN3CCN (C4=CC=CC (Cl) =C4Cl) CC3) C=C2N1”. The input for the model is the “one-hot” representation of a SMILES string, where each string is split into tokens. In this case, the first token is “O”, which is converted into a “one-hot” vector and is input into the language model. Then, the model updates its hidden state and outputs the probability distribution over the next possible tokens, which in this example is decoded as “=“. Feeding the one-hot vector of “=” to the model causes the hidden state to update in the next iteration, resulting in the next token. This iterative token-by-token procedure continues until the character “\n” is encountered, which is then added to denote the end of the SMILES string, ultimately affording the SMILES for aripiprazole. To ensure that the RNN can generate rational, drug-like molecules, the model was first trained with 3.1 million drug-like molecules from the *ZINC* database. The model converged quickly (within 3 epochs) (Figure S2) due to the size of the training data set.

We sampled 3.3 million SMILES strings based on the well-trained RNN model, and of these, more than 95% were valid and chemically feasible molecules. A validity check was performed by parsing the generated SMILES strings with the RDKit toolkit. The repetition rate between the generated molecules and the training set was only 0.21%. Random examples of the generated molecules are depicted in Figure S3.

Subsequently, we investigated whether the generator learned the distribution of the molecular properties represented in the training set. Several physicochemical properties of the generated compounds were calculated, these include the molecular weight (MW), BertzCT, rotatable bonds, Wildman-Crippen partition coefficient (logP), total polar surface area (TPSA) ^31^, and number of H-acceptors and H-donors. These properties were calculated using the RDKit with a set of 10000 molecules randomly selected from both the training set and the generated set. To better demonstrate the distribution of structural and physicochemical properties of the generated compounds, t-distributed stochastic neighbor embedding (t-SNE) was conducted for dimensionality reduction^32^. In addition to visualizing the distribution of physicochemical properties, a t-SNE plot of the extended connectivity fingerprint 4 (ECFP4) fingerprint was prepared to visualize the chemical space covered by the generated samples and the samples in the training set. As shown in Figure 1C, D, the distributions of the generated and the training set compounds overlapped significantly, indicating that the generated samples well represent the properties of the samples training set.

Previous studies have shown that the presence of heterocyclic or fused aromatic rings in a compound may contribute to their interactions with the receptor and the efficacy of the compound as an antipsychotic drug^33^. Thus, the numbers of three types of ring systems, namely, heterocyclic, aromatic and aliphatic rings, were counted for both the designed molecules and the molecules in the *ZINC* set (Figure 1E). Synthetic accessibility (SA) and drug-likeness are important considerations in the development of novel drug candidates. In this study, synthetic accessibility was evaluated using the SA score based on a combination of fragment contributions and a complexity penalty^34^. This method characterizes SA as a score between 1 (easy to make) and 10 (very difficult to make). Drug-likeness was assessed based on the quantitative estimate of drug-likeness (QED) index, which measures the consistency of a given compound with currently known drugs in terms of structural and physicochemical properties^35-36^. Then, the SA and QED scores for both the training set and the generated set (10000 molecules randomly sampled from each set) were calculated. The results (Figure 1E, Figure S4A) illustrated that the distributions of these properties in the generated molecules highly resembles that of the molecules in the *ZINC* data set. In addition, most of the de novo molecules possess good QED and SA scores.

### 2.2. Generating the Molecular Library Focused on D_1_/D_2_/5-HT_1A_/5-HT_2A_ Receptors

We aimed to establish a library of compounds to simultaneously target D_1_, D_2_, 5- HT_1A_ and 5-HT_2A_ receptors. To compile the training data set, molecules reported having pIC_50_ > 6 or pEC_50_ > 6 for D_1_/D_2_/5-HT_1A_/5-HT_2A_ receptors were collected from the GLASS, Reaxys and SciFinder databases. In total, 10286 molecules (580793 symbols) were obtained to fine-tune the pretrained RNN model. The fine-tuned model converged after 7 epochs, which is slower than the pretrained model (Figure S2). Then, 10,000 SMILES sequences from the fine-tuned generative model were sampled, and the same investigation and comparison were conducted on the fine-tuning set using the properties mentioned above (Figure 1C-E). As shown in Figure 1F, G, the generated molecules possess similar physicochemical properties and occupy the same chemical space as the fine-tuning set. Unlike the molecules generated by the pretrained model (Figure 1E), the molecules generated with the fine-tuned model possess higher proportions of heterocyclic, aromatic and aliphatic rings, which is consistent with the fine-tuning set (Figure 1H). The distributions of QED and SA scores among the generated molecules is also similar to that of the compounds in the fine-tuning set (Figure 1H, Figure S4B).

The structures of a random sample of molecules from the fine-tuned model are depicted in Figure S4C. As expected, known antipsychotic drugs such as aripiprazole and risperidone can be reproduced by the fine-tuned model, suggesting that the model has learned the structural distribution of published antipsychotic drugs. Molecules similar to the published D_1_/D_2_/5-HT_1A_/5-HT_2A_ modulators were also generated, indicating that the model could generate close analogs of molecules in the fine-tuning set. Molecules that were not as closely related to the training set were also produced, which showed that the fine-tuned model could provide new chemotypes or scaffolds for drug optimization. The similarity of each compound was calculated by a nearest-neighbor similarity analysis using an ECFP4-based similarity method with the Tanimoto index^37^. A larger similarity value indicated that the generated molecule was more similar to the training set.

However, under iterative training on the same training set, a high proportion of the molecules produced by the generative model will be highly similar to those in the training set. To address this issue, a discriminative model to evaluate whether the generated molecule is active needs to be incorporated; this model will be explained in the next section. The top-ranked molecules with high predicted activities were used to expand the fine-tuning set in the next iteration. These steps were repeated to favor the generation of molecules with high activity and low similarity.

### 2.3. Deep discriminative model for activity prediction by MTDNN

MTDNNs can simultaneously model more than one task, which facilitates the reuse of features that are learned from multiple tasks and share statistical strength. Here, multitask DNNs were employed to establish our predictive model for developing novel multitarget antipsychotic drugs that act on D_2_/5-HT_1A_/5-HT_2A_ receptors. Since it has been demonstrated that the multitask performance would be improved by incorporating more relevant tasks, the dopamine D_1_ receptor was also considered due to its high degree of structural homology to D_2_ and the large amount of bioactivity data available. Accordingly, regression models for predicting the pIC_50_ and pEC_50_ values against D_1_/D_2_/5-HT_1A_/5- HT_2A_ receptors were established (Figure 2A). A transfer learning strategy was then employed to further improve the performance of the discriminative model, which focuses on storing knowledge gained while solving one problem and applying it to a different but related problem. As shown in Figure 2A, the aim of transfer learning in this case is to learn general features from a larger data set and apply them to a smaller data regime by fine-tuning. Herein, a multitask model was first trained with the K_i_ data sets, and the learned weights from the pK_i_ multitask DNN model were then used to initialize the pIC_50_ and pEC_50_ multitask DNN models.

**Figure 2.**
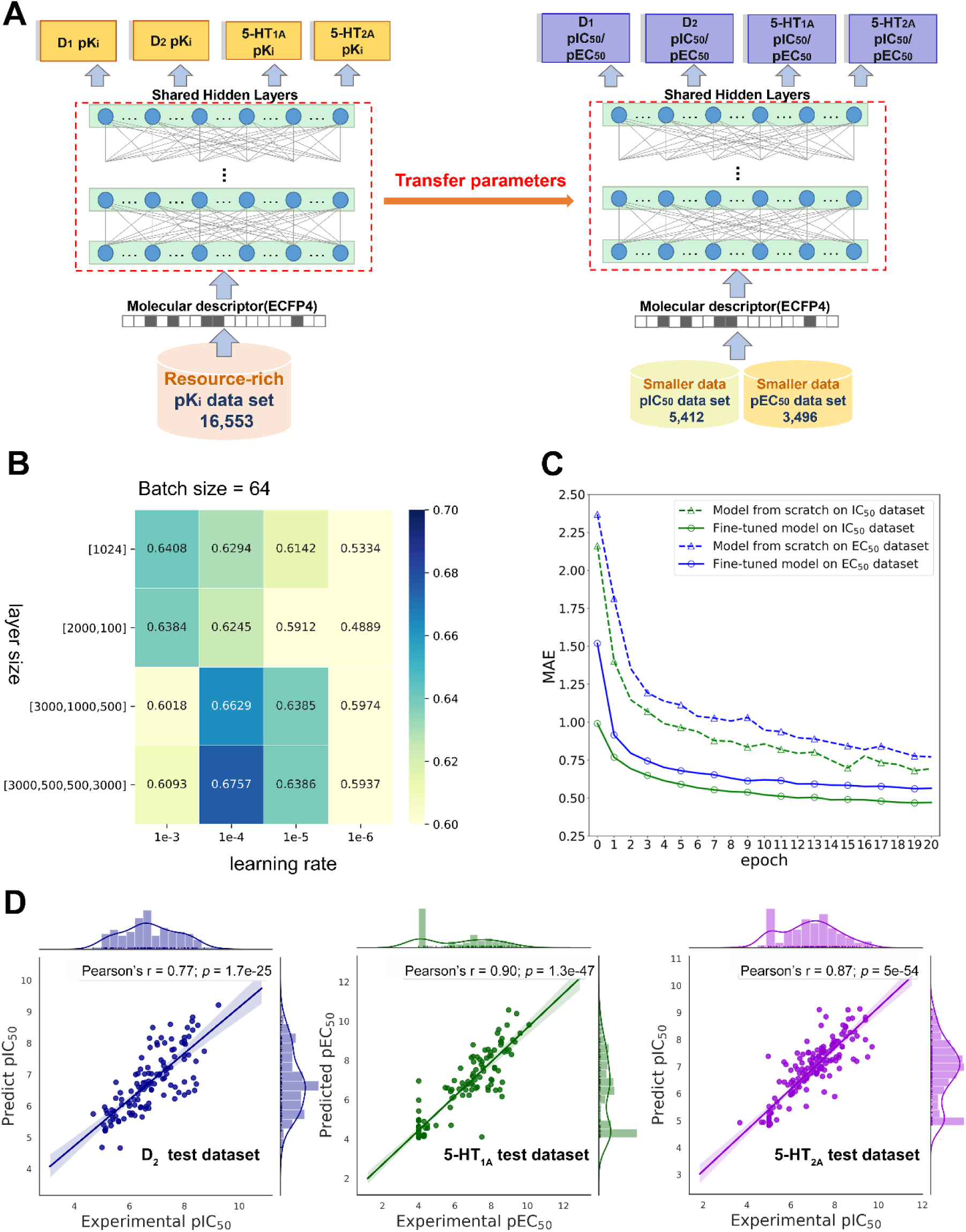
Deep discriminative model for activity prediction by MTDNN. (A) Architecture of the transfer learning from the larger data set (pK_i_ data set) to the smaller data set (pIC_50_, pEC_50_ data set). The MTDNN model takes ECFP4 as input descriptor and produces predicted values of p-bioactivity (pK_i_, pIC_50_ and pEC_50_). (B) The performance results of the MTDNN model on pK_i_ data set with different hyperparameters. The annotation marked in each block indicate the r^2^ of the test data set. The darkest color corresponds to the best performance model. (C) MAE comparison for the IC_50_ test data set and EC_50_ test data set over 20 epochs, where the green lines indicate the MTDNN model trained with the IC_50_ test data set, the blue lines indicate the MTDNN model trained with the EC_50_ test data set. The lines containing triangle markers correspond to randomly initialized models, and the lines containing circle markers correspond to the MTDNN model trained with weights learned from the MTDNN model pretrained on the pK_i_ data set. (D) Correlation between the experimental values and the corresponding predicted values on the test data sets of D_2_, 5-HT_1A_ and 5-HT_2A_, respectively. Upper and right column bar graphs show the distributions of experimental and predicted values. The Pearson’s r value and *p* value for each scatter plot are shown in the upper area.

A standard grid searching method was used to determine the hyperparameters, the predictive performance on the pK_i_ test data set of the MTDNN models is shown in Figure 2B. With the learning rate of 1e-4, the layer size of [3000, 500, 500, 3000], and the batch size of 64 on the pK_i_ data set, the model achieved best performance with an r^2^ of 0.68 and a MAE of 0.45. Next, we adopted the parameters of this best performance pK_i_ model and transferred the parameters to the MTDNN models on the pIC_50_ and pEC_50_ data sets for 100 epochs. As depicted in Figure 2C, the fine-tuned model converged more quickly and showed a lower MAE than did the randomly initialized model. By exploiting and identifying the common representations between individual properties, transfer learning increased the accuracy of the models and significantly improved the efficiency of the training process. Finally, we obtained the test model accuracy expresses as r^2^ of 0.71, 0.71 and MAE is 0.47 and 0.54 respectively for IC_50_ multitask model and EC_50_ multitask model. Then, the Pearson’s correlation coefficient between experimental values and the predicted values on D_2_, 5-HT_1A_ and 5-HT_2A_ test data set were calculated. As shown in Figure 2D, there’s a stronger linear correlation between the experimental and predicted results, with Pearson’s r values of 0.77, 0.90 and 0.87 for D_2_, 5-HT_1A_ and 5-HT_2A_ test data set respectively.

### 2.4. Adaptive antipsychotic design

To generate molecules that exhibited high activity and low similarity, the generative model was combined with a discriminative model. The molecules generated in each round of retraining were virtually evaluated by the MTDNN model and further screened based on the other filter criteria, including QED, SA, and structural novelty. A compound will be filtered out if its QED score is less than 0.5, its SA score is greater than 5, or it has a long aliphatic chain with more than 8 carbon atoms. The compounds that met these criteria were then collected, and the prioritized structures with higher predicted activities (mean predicted values for the D_2_/5-HT_1A_/5-HT_2A_ receptors) were resampled to enrich the fine-tuning set for the next iteration. Furthermore, the average activity threshold for the discriminative model was progressively increased to select molecules with higher predicted activities values for the D_2_, 5-HT_1A_ and 5-HT_2A_ receptors. The initial threshold value was set to 6, and after the second epoch, it was increased by 0.1 after each epoch until the generative model converged.

A total of 10286 molecules with reported activities toward dopamine and serotonin receptors were collected, and they were used to fine-tune the pretrained generative model. Virtual screening and fine-tuning of the model with high-ranking generated molecules were carried out for 20 epochs. As shown in Figure 3A, the loss plateaued after the 14-th epoch. Notably, after 2 epochs, more than 50% of the molecules produced by the model were active, and the proportion quickly converged after 8 epochs. To explore how the predicted activity changed during the iterative process, molecules with predicted average activities greater than the activity threshold were collected from each epoch. As shown in Figure 3B, the predicted average activity values monotonically increased and began to plateau after the 14-th epoch.

**Figure 3.**
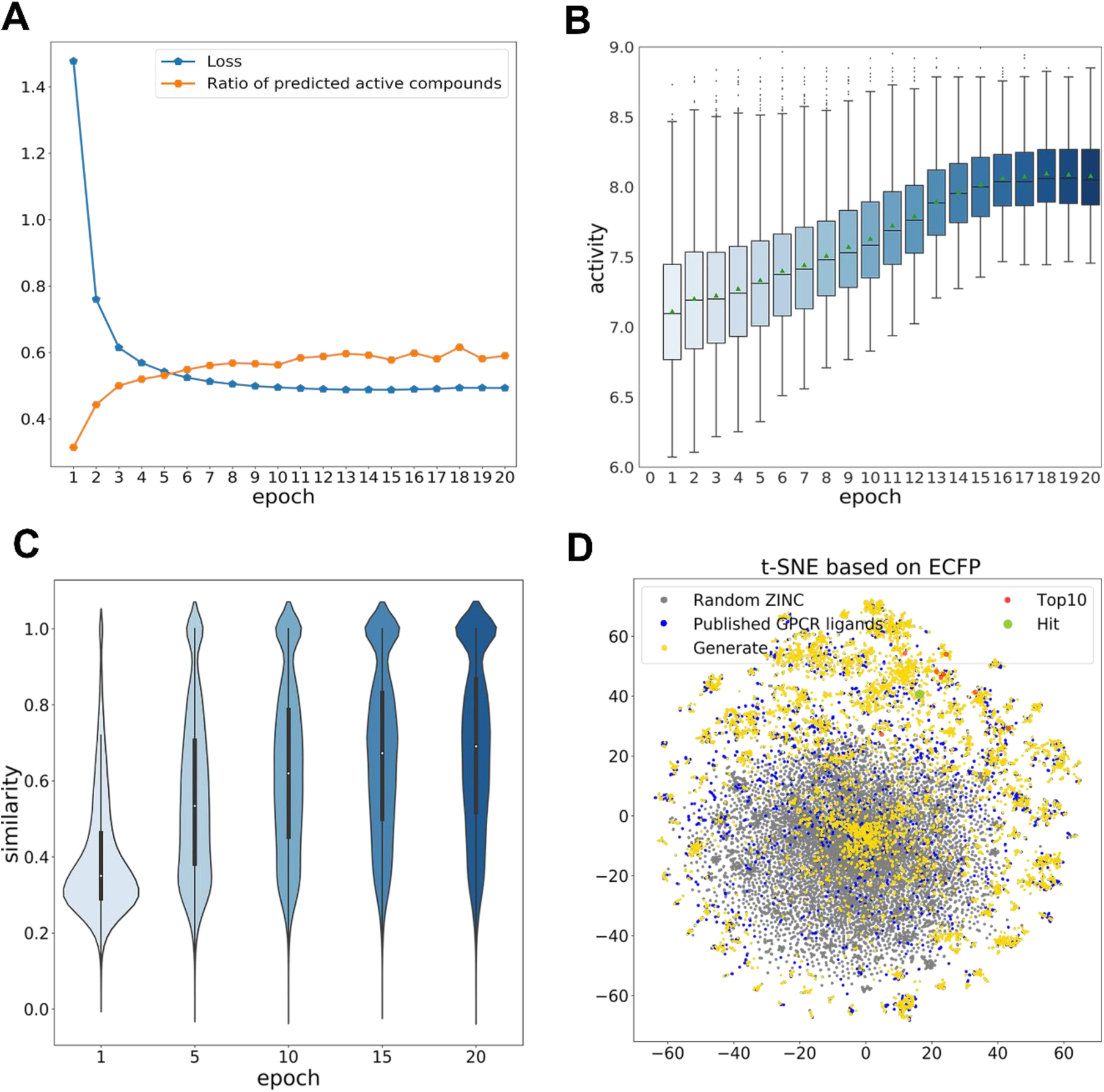
Adaptive antipsychotic design process and analysis of the physicochemical properties of the generated molecules. **(A)** Loss curve of the fine-tuned RNN model and the proportion of predicted active molecules during the fine-tuning process with the GPCR data set over 20 epochs. **(B)** Changes in the predicted activity values on the D_2_/5-HT_1A_ / 5-HT_2A_ receptors with epochs of fine-tuning. **(C)** Violin plot of the nearest-neighbor ECFP4-Tanimoto similarity distribution of the generated molecules relative to the training set at epochs 1, 5, 10, 15, and 20. **(D)** Chemical space analysis by multidimensional scaling on t-SNE. The colored dots represent random samples of the pretraining data (gray), the fine-tuning set (blue), molecules generated with the fine-tuned model (gold), the top 10 de novo designs (red), and hits (green). The retrained generator populates the chemical space around the training molecules.

Finally, 10000 molecules were sampled from the 14-th generative model. To assess the novelty of the de novo molecules generated by the joint system, a similarity analysis was performed. The violin plot in Figure 3C shows the distribution of the Tanimoto index values of the generated molecules and their nearest neighbors in the training set at 1, 5, 10, 15 and 20 epochs. Notably, the model started to output an increasing number of molecules similar to the target-specific training set after 5 epochs. The distribution suggests that the model has learned to generate close analogs but can also generate novel molecules that are not highly similar to the fine-tuning set.

To understand how the generative model populates the chemical space with new molecules, a visualization based on t-SNE by multidimensional scaling was performed using the ECFP4 fingerprints. The colored dots represent random samples of the pretraining data (gray), published D_2_/5-HT_1A_/5-HT_2A_ modulators (blue), and molecules generated with the fine-tuned model (gold). As shown in Figure 3D, the newly generated molecules populate the chemical space of the pretraining set—both within the fine-tuning set and in the adjacent empty space. The results demonstrate the ability of the generative model to produce novel chemical entities within the training data set domain, which is consistent with what has been shown previously^38-39^.

To further understand the physicochemical properties of the generated molecules, several molecular descriptors were calculated: MolWt, clogP, TPSA, etc. In addition, instead of using Lipinski’s rule of five (RO5) as a selection criterion, QED and SA were adopted, which impose stricter limitations on the molecules^40^. Figure S5 shows that the generated molecules reproduced the same distribution as the training molecules that have been reported to be active. Overall, these results suggested that the joint system can produce structures suitable for organic synthesis and subsequent biological evaluation.

### 2.5. Hit Selection and Assay Verification

As mentioned above, a focused library of D_2_, 5-HT_1A_ and 5-HT_2A_ receptors was designed and virtually optimized by the combined automated system. Previous studies on psychotropic drugs have suggested that a balanced activity profile against dopamine and serotonin receptors is important for minimizing side effects^41-42^. To retain the potency and avoid undesired selectivity for the three targets among the generated molecules, two primary selection criteria were set: (a) high potency for the D_2_, 5-HT_1A_ and 5-HT_2A_ receptors and (b) a balanced activity profile, meaning that the potency ratio between any two receptors should be no greater than 10, and the potency ratio between 5-HT_1A_R/D_2_R should greater than 1.0. Moreover, to obtain novel molecules, we kept only the molecules that showed a Tanimoto similarity no greater than 0.75 relative to the training set molecules. Starting with the 10, 000 molecules generated from the 14-th epoch model, the pan-assay interference compounds (PAINS) were removed by pipeline Pilot^43^, then we filtered the molecules based on the above criteria, and the retained molecules were sorted according to their predicted average activities against the D_2_, 5-HT_1A_ and 5-HT_2A_ receptors. An ideal property profile for the design of high-quality central nervous system (CNS) drugs has been proposed^44^. These parameters include clogP in the range 2–5, TPSA less than 76 Å^2^, log BB greater than –1 and positive QikProp. When integrating the QikProp properties into the multiobjective optimization, the complexity of the calculations may lead to a slower generating speed and less flexibility. Therefore, the QikProp properties of the candidates were calculated separately. The top 5 molecules and their predicted properties are shown in Table 1.

**Table 1.**
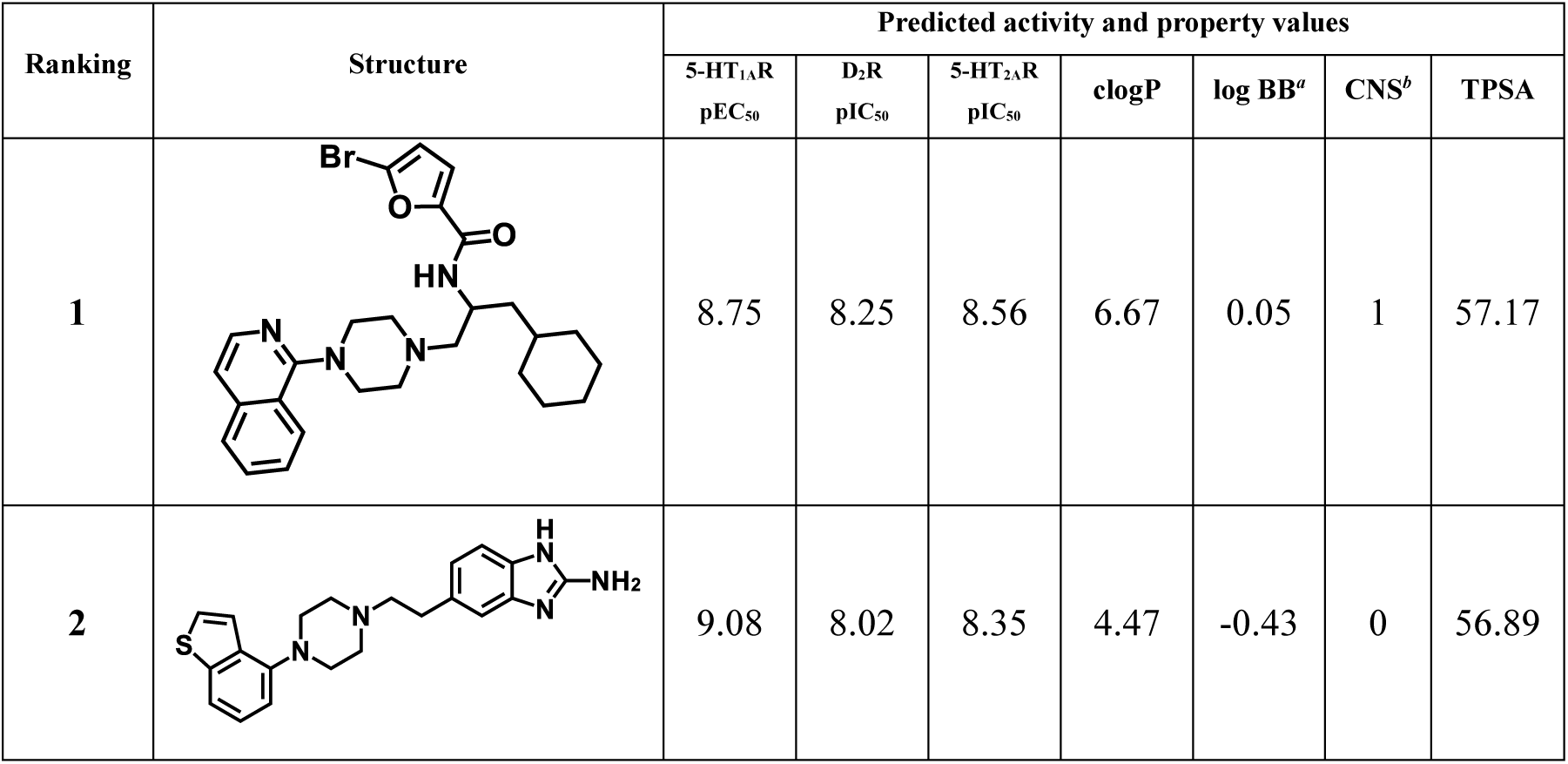
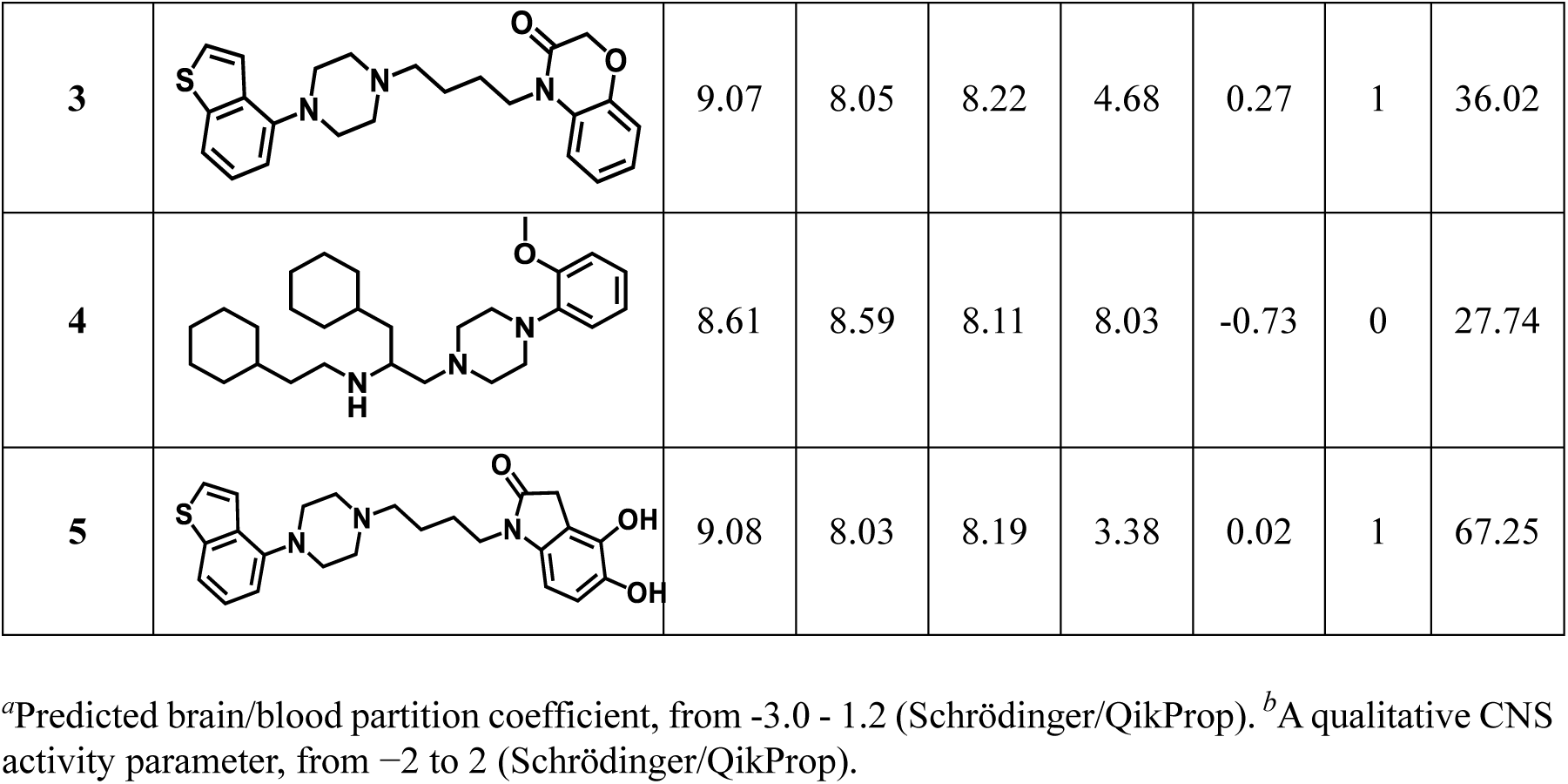
Top 5 molecules generated by the joint deep learning system

| Ranking | Structure | Predicted activity and property values |  |  |  |  |  |  |
| --- | --- | --- | --- | --- | --- | --- | --- | --- |
|  |  | 5-HT<sub>1A</sub>R<br>pEC<sub>50</sub> | D<sub>2</sub>R<br>pIC<sub>50</sub> | 5-HT<sub>2A</sub>R<br>pIC<sub>50</sub> | clogP | log BB<sup>a</sup> | CNS<sup>b</sup> | TPSA |
| 1 |  | 8.75 | 8.25 | 8.56 | 6.67 | 0.05 | 1 | 57.17 |
| 2 |  | 9.08 | 8.02 | 8.35 | 4.47 | -0.43 | 0 | 56.89 |
| 3 |  | 9.07 | 8.05 | 8.22 | 4.68 | 0.27 | 1 | 36.02 |
| 4 |  | 8.61 | 8.59 | 8.11 | 8.03 | -0.73 | 0 | 27.74 |
| 5 |  | 9.08 | 8.03 | 8.19 | 3.38 | 0.02 | 1 | 67.25 |

<sup>a</sup>Predicted brain/blood partition coefficient, from -3.0 - 1.2 (Schrödinger/QikProp). <sup>b</sup>A qualitative CNS activity parameter, from -2 to 2 (Schrödinger/QikProp).

Compounds **1** and **4** have clogP values>5.0, indicating poor oral absorption and intestinal permeability^45^. Furthermore, in our experience, introducing bulky substituents such as a cyclohexylmethyl group on the middle linker of the arylpiperazine derivative is detrimental for DA and 5-HT receptor binding. Except for **1** and **4**, the other compounds have clogP and log BB values within the desirable ranges, suggesting a reasonable probability that they will be able to penetrate the blood-brain barrier. The presence of two phenolic hydroxyl groups in the structure of **5** may contribute to its biological activity but could lead to a poor PK profile; therefore, **5** was also excluded from our synthetic plan along with compounds **1** and **4**. Starting from compound **2**, a series of arylpiprazine derivatives with a ethyl linker were synthesized and evaluated as potential antipsychotic drugs, and this part of our study will be discussed in the near future. Herein, we discuss the synthesis and results of compound **3**.

The synthesis route for **3** was illustrated in Scheme 1. Commercial **3-a** was alkylated by 1,4-dibromobutane by treatment with NaH in *N,N*-dimethyl formamide (DMF) to afford intermediate **3-b**. The synthesis of compound **3** was accomplished by coupling **3-b** with 1-(benzo[b]thiophen-4-yl) piperazine hydrochloride in the presence of K_2_CO_3_ in MeCN.

Then, functional activity assays on intracellular cAMP levels were employed to determine the activities of compound **3** on D_2_R, 5-HT_1A_R and 5-HT_2A_R. Compound **3** exhibited good activities against D_2_R and 5-HT_2A_R with IC_50_ values of 216 nM and 1.64 nM, respectively, and showed good agonist activity on 5-HT_1A_R with an EC_50_ value of 0.51 nM, which is in accordance with the activities predicted by MTDNN (Table 1). As shown in Figure 3D, compound **3** and the top 10 molecules populate the same chemical space as known antipsychotic drugs. Encouraged by the good activity profile of compound **3**, it was chosen as the hit compound for further structural optimization to identify more favorable molecules.

**Scheme 1.**
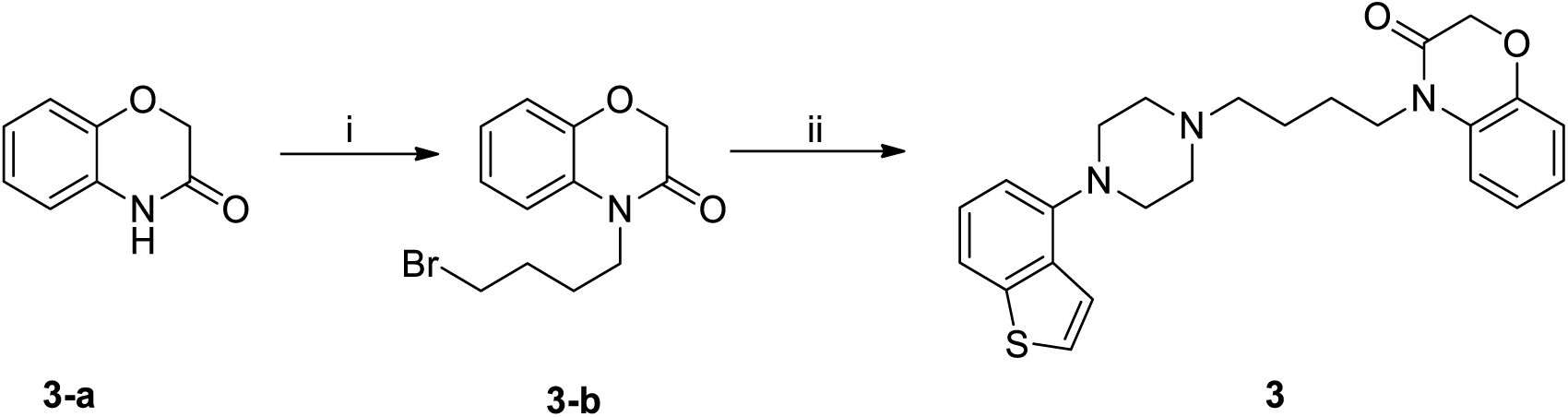
Reagents and conditions: (i) 1,4-dibromobutane, NaH, DMF, 30 °C; (ii) 1- (benzo[b]thiophen-4-yl)piperazine hydrochloride, K_2_CO_3_, MeCN, reflux.

### 2.6. Hit expansion and in vitro Pharmacological Profile Determination

Considering compound **3** as a hit, 10 analogs were designed by introducing different linkers or heterocyclic moieties. To improve efficiency and avoid unnecessary time and labor costs, the MTDNN models were used for the preliminary screening. The predicted activity profiles of the 10 analogs (compounds **12**-**15** showed in Table S1) indicated that their predicted average functional activities against the three targets gradually increased as the linker length increased from two to four carbons, which was consistent with previous studies. Previous SAR studies on psychotropic drugs also demonstrated that linker length plays an important role in determining receptor function and that the a butylene is generally considered to have the optimal length^46-48^. Thus, an investigation of the linker moiety focused on chains approximately four carbons long. After excluding the 4 analogs that showed low predicted activities against the D_2_/5-HT_1A_/5-HT_2A_ receptors (Table S1), the remaining 6 analogs were synthesized for further biological investigations following procedures similar to that described for **3** (Schemes 2 and 3). The details of the syntheses of the 6 analogs are described in the Methods section.

**Scheme 2.**
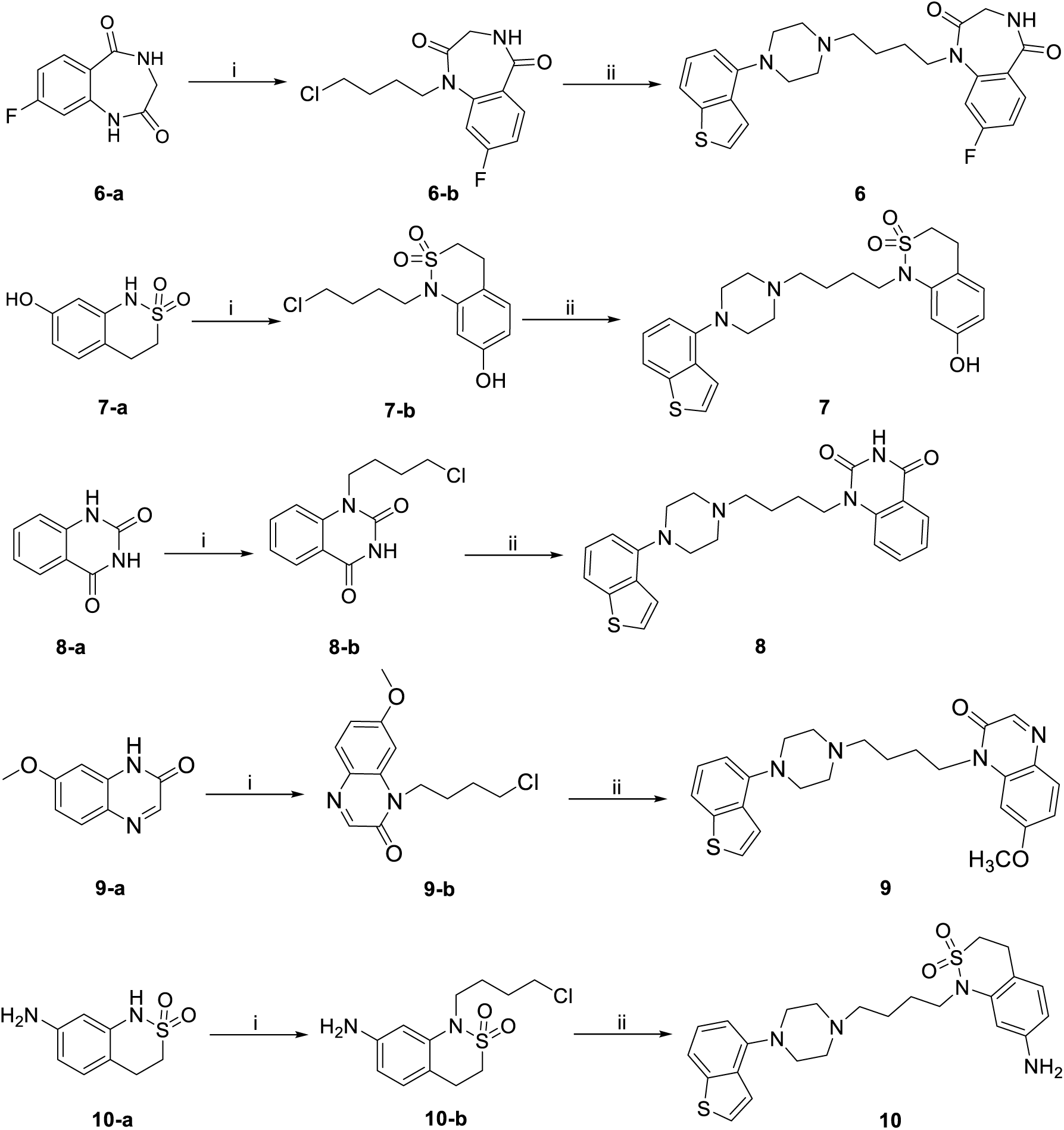
Reagents and conditions: (i) 1-bromo-4-chlorobutane, K_2_CO_3_, DMF, rt; (ii) 1- (benzo[b]thiophen-4-yl)piperazine hydrochloride, K_2_CO_3_, MeCN, reflux.

**Scheme 3.**
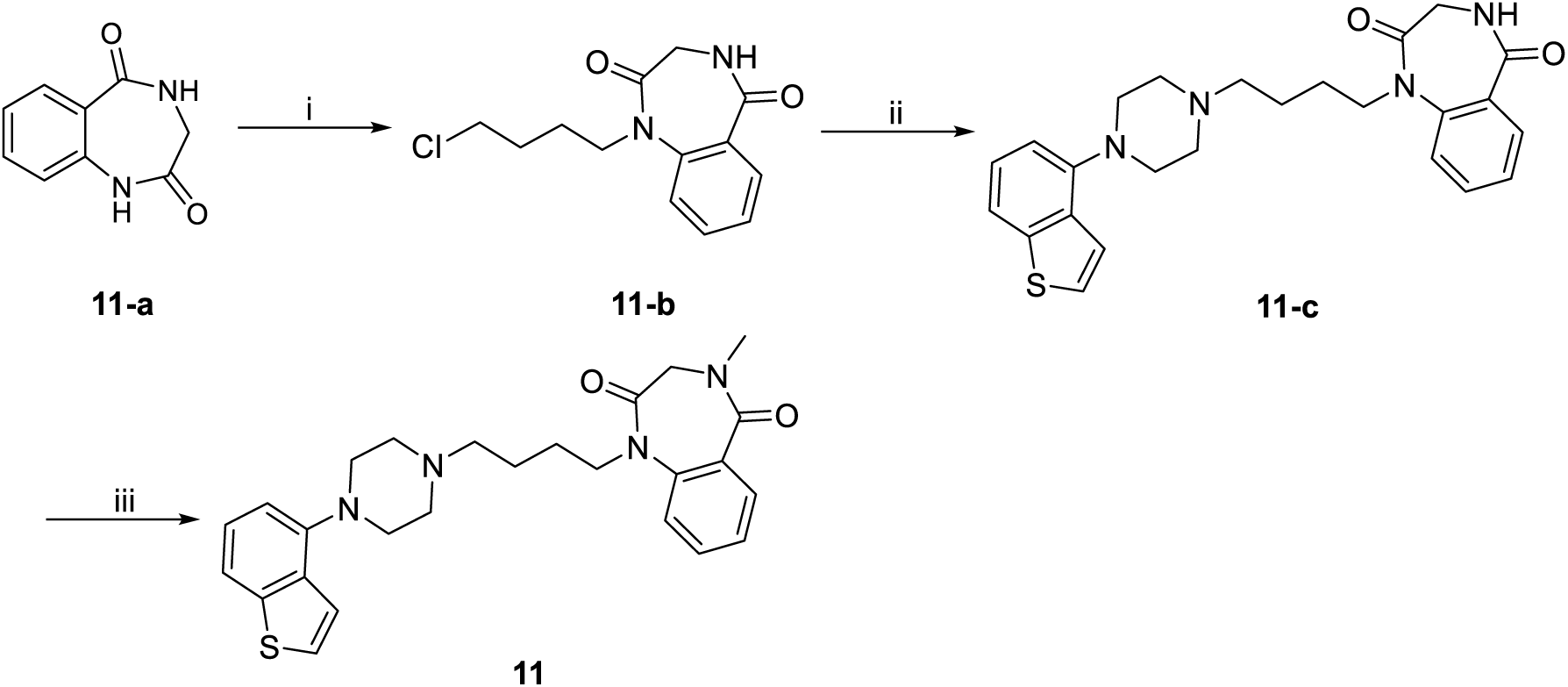
Reagents and conditions: (i) 1-bromo-4-chlorobutane, K_2_CO_3_, DMF, rt; (ii) 1- (benzo[b]thiophen-4-yl)piperazine hydrochloride, K_2_CO_3_, MeCN, reflux; (iii) NaH, Me_2_SO_4_, DMF, rt.

Then, the activity profiles of the 6 analogs against the D_2_/5-HT_1A_/5-HT_2A_ receptors were characterized *in vitro*. As shown in Figure 4A, the results showed that the designed compounds exhibited potent potency toward D_2_/5-HT_1A_/5-HT_2A_ receptors, which was consistent with the predictive models. The present “atypical” antipsychotics are considered to possess potent D_2_ and 5-HT_2A_ receptor affinity but relative lower 5HT_1A_ affinity. Considering that compounds of stronger affinity toward 5HT_1A_ receptors have rarely been systematically studied, we developed this workflow and designed several compounds that meet our expectations in a shorter period of time. It was observed that these compounds exhibited potent agonist activity against 5-HT_1A_R, with a stronger potency than D_2_R, and these compounds can be used as molecular probes in the exploration of the augmentation strategy in treatment of schizophrenia. In agreement with the predictive models, compared with compounds **11**, compound **6** bearing a fluorine moiety displayed higher potency on 5-HT_1A_ agonism activity. The MTDNN predicter promoted the optimization process of the hit compound to meet the initial expectations, and it can be used as an efficient approach to study the corresponding structure-activity relationship.

**Figure 4.**
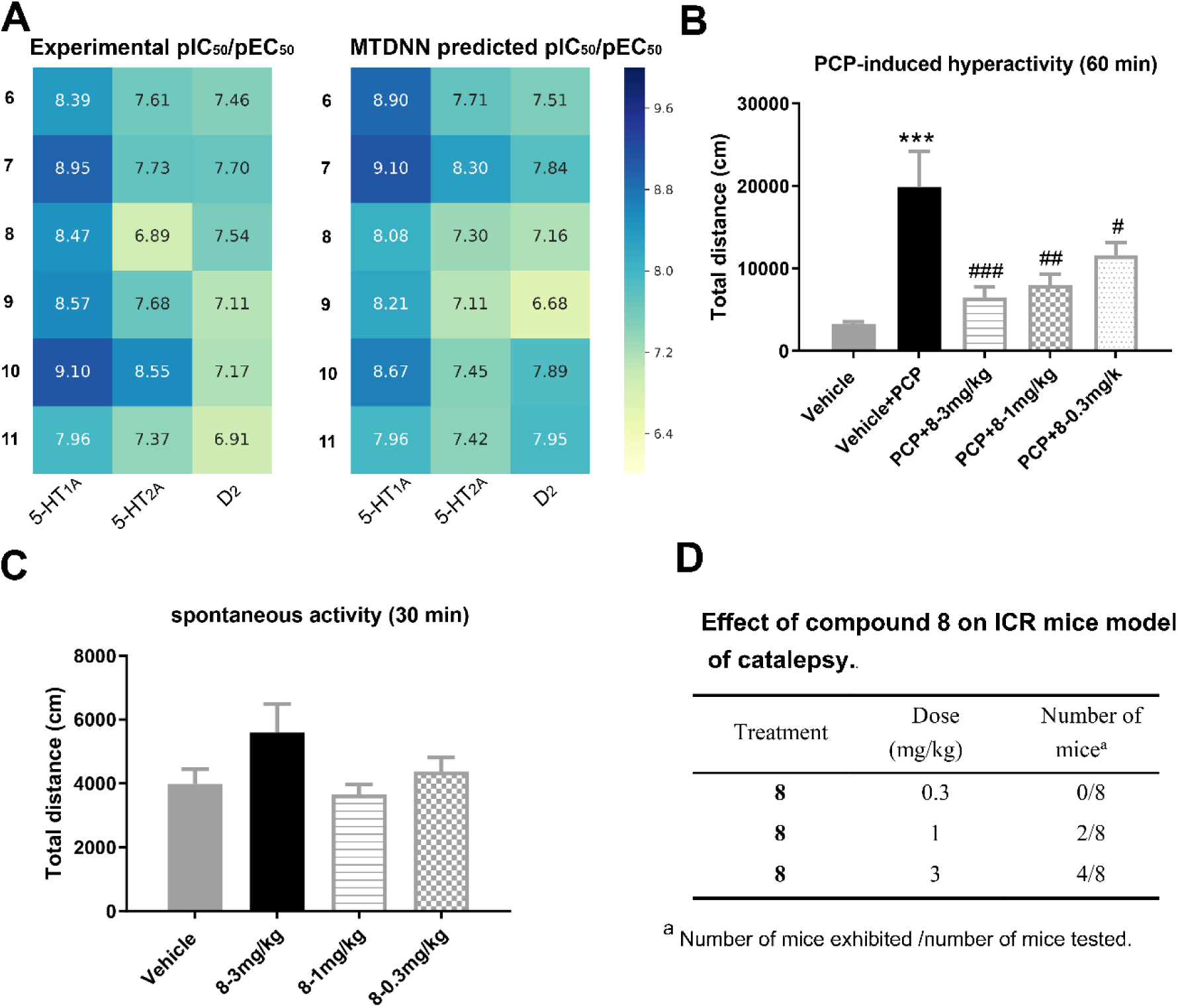
*In vitro* pharmacological profile determination of 6 analogs of compound 3 and *in vivo* behavioral evaluation of compounds **8**. (A) Comparison of the polypharmacology profiles for the MTDNN model score and *in vitro* experimental activity profiles of the 6 analogs against the D_2_/5- HT_1A_/5-HT_2A_ receptors. (B) Effects of compound **8** administered po on PCP-induced hyperactivity in mice (PCP: 7 mg/kg, ip). Locomotor activities were measured for 60 min following PCP administration, and the results are expressed as the mean ± SEM of the distance traveled. (C) Effects of compound **8** administered po on spontaneous activity in mice. The locomotor activities were measured during the 30 min following drug administration, and the results are expressed as the mean ± SEM of distance traveled. Statistical evaluation was performed by one-way ANOVA followed by Dunnett’s post hoc test. #*p* < 0.05 versus PCP treatment; ##*p* < 0.01 versus PCP treatment; \*\*\**p* < 0.001 versus vehicle treatment. (D) Effect of compound **8** on ICR mice model of catalepsy.

### 2.7. In vivo behavioral evaluation of the selected compounds

Based on the *in vitro* pharmacological profiles, compound **3** and these 6 analogs were then subjected to *in vivo* behavioral studies to verify their effects on schizophrenia Antipsychotic efficacy is usually assessed by the phencyclidine (PCP)-induced hyperactivity model^49-50^. PCP, an N-methyl-D-aspartic acid receptor (NMDAR) antagonist, has been found to simulate schizophrenia^51^, and in rodents, this effect can be reversed by antipsychotic drugs^52^. Then the antipsychotic activities of the above 7 compounds were studied in the PCP-induced locomotor hyperactivity test in ICR mice (Figure S6). In this model, compound **8** produced more potent efficacy than other compounds and aripiprazole at 3 mg/kg (Table 2 and Figure S6).

**Table 2.**
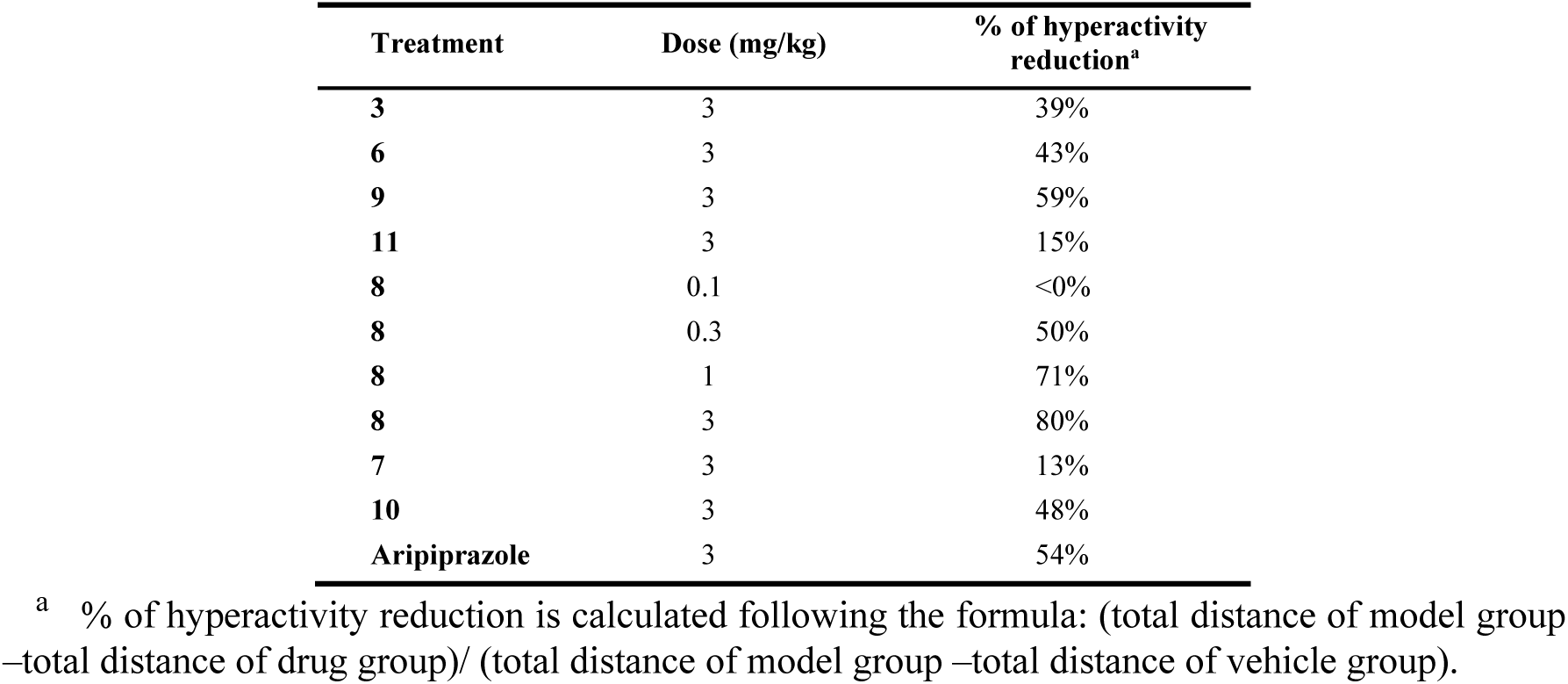
Effect of compound **3** and 6 analogs on ICR mice model of PCP-induced hyperactivity response.

Besides, it is worth to note that compound **8** reversed the PCP-induced hyperactivity dose-dependently with the minimum effective dosage ranging between 0.1 and 0.3 mg/kg (Figure 4B and Figure S6A). Moreover, as the spontaneous locomotor activity of mice after administration of compound **8** at effective doses (0.3 to 3 mg/kg) was not influenced (Figure 4C), its antipsychotic-like effects should be specific. In contrast, although aripiprazole could inhibit PCP-induced hyperactivity at 3 mg/kg (Figure S6D), the spontaneous locomotor activity of mice at this dosage was significantly influenced, suggesting a potential side effect of sedation.

Catalepsy is a frequently used model to predict liability of antipsychotics to induce EPS side effect in humans. As shown in Table 3, Compound **8** did not cause remarkable catalepsy in mice at the dosage of 1 mg/kg, indicating an acceptable safety profile.

**Table 3.** Detailed information of the modeling data sets

| UniProt<br>ID | Gene<br>Name | Protein Name | Number of compounds with<br>different bioactivity types |  |  |
| --- | --- | --- | --- | --- | --- |
| | | | $K_i$ | $IC_{50}$ | $EC_{50}$ |
| P21728 | D<sub>1</sub> | D (1) dopamine receptor | 1330 | 216 | 636 |
| P14416 | D<sub>2</sub> | D (2) dopamine receptor | 6121 | 1470 | 649 |
| P08908 | 5-HT<sub>1A</sub> | 5-hydroxytryptamine receptor 1A | 5228 | 1710 | 1347 |
| P28223 | 5-HT<sub>2A</sub> | 5-hydroxytryptamine receptor 2A | 3874 | 2016 | 864 |

## 3. Conclusions

In this work, a new workflow for the automated design of de novo antipsychotic molecules was developed based on deep neural networks. An efficient system composed of an RNN generative model and an MTDNN discriminative model was established to create molecules with a specific multitarget profile. Several iterations with this system resulted in the identification of high-scoring compound **3**, which exhibited potent activities against D_2_, 5-HT_1A_ and 5-HT_2A_ receptors in biochemical experiments. Taking compound **3** as a hit, a simple structural-activity relationships (SAR) experiment was performed using the MTDNN model, and 6 analogs of compound **3** identified in this study were synthesized. The bioactivities of these six compounds on the D_2_/5-HT_1A_/5-HT_2A_ receptors were evaluated. Among these compounds, compound **8** exhibited promising activities on the D_2_, 5-HT_1A_ and 5-HT_2A_ receptors and were demonstrated to exhibit potent antipsychotic-like effects in behavioral studies with low potential for sedation and catalepsy. Finally, compound **8** showed promise for development as novel antipsychotic drugs.

In conjunction with wet lab experiments, this study substantiated the practicality and feasibility of using AI in de novo drug design. We focused on designing de novo antipsychotic drugs as a convenient test case and developed an automated drug design workflow for achieving multitarget profiles. Overall, the workflow is applicable to all drug-target classes when sufficient structure-activity data exist to create discriminative models. The proposed method generates efficient tools for designing small molecules with a specific multitargeting profile. This work could also be extended to achieve selectivity over other undesired drug targets. Additionally, the accuracy of the predictive model might be improved by training the model with more useful information, such as protein structure and protein sequences, as it becomes available, and this will be explored in the near future.

## 4. Methods and experimental

### 4.1. RNN modeling data and curation

For the pretraining set, 310703 SMILES strings of drug-like molecules were collected from the *ZINC* database. The SMILES strings of the molecules were canonicalized by the RDKit toolkit and transformed into sequences of symbols found in the training set.

The fine-tuning sets were obtained from GLASS, Reaxys and SciFinder databases. First, molecules from the DNN modeling data set (collected from GLASS and Reaxys databases) were screened, and molecules with pIC_50_ or pEC_50_ values greater than 6 that are active against any of the D_2_/5-HT_1A_/5-HT_2A_ receptors were collected. To expand the multitarget training set, molecules from SciFinder that met the above criteria were added. Finally, 10286 molecules were obtained, all of which were canonicalized with the RDKit toolkit.

### 4.2. Recurrent neural network

RNNs are artificial neural networks that are able to use their internal state (memory) to process sequences of inputs. The RNN architecture unfolds over time (as shown in Figure 5A), which allows RNNs to process sequences of input vectors *s*_1:*n*_ = (*s*_1_, *s*_2_, …, *s*_*n*_). The RNN model is trained by maximum likelihood estimation. The model centers on learning a probability distribution over all the next possible symbols (symbols vocabulary) and aims to maximize the likelihood assigned to the correct token. The learned probability distribution of a sequence s can be written as follows:

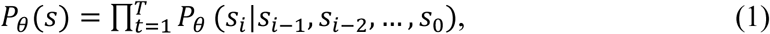

where the parameters *θ* are learned from the training set and *s*_*i*_ is the *i*-th symbol at step *t*_*i*_ ∈ *T*. The RNN model takes a sequence of input vectors *s*_1:*n*_ = (*s*_1_, *s*_2_, …, *s*_*n*_), and an initial state vector *h*_0_, and the hidden state updates and returns a sequence of output vectors *o*_1:*n*_ = (*o*_1_, *o*_2_, …, *o*_*n*_). At each time step *t*, the hidden state *h*_*t*_ is updated as follows:

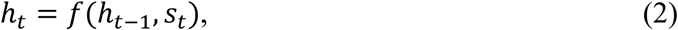

where *f* is a recursive activation function. *f* may be as simple as a sigmoid function or as complex as a LSTM unit, which is a special kind of RNN that is capable of learning-term dependency. The performance of the RNN was greatly improved by the use of microarchitectures as LSTM cells, avoiding the long-term dependency problem.

**Figure 5.**
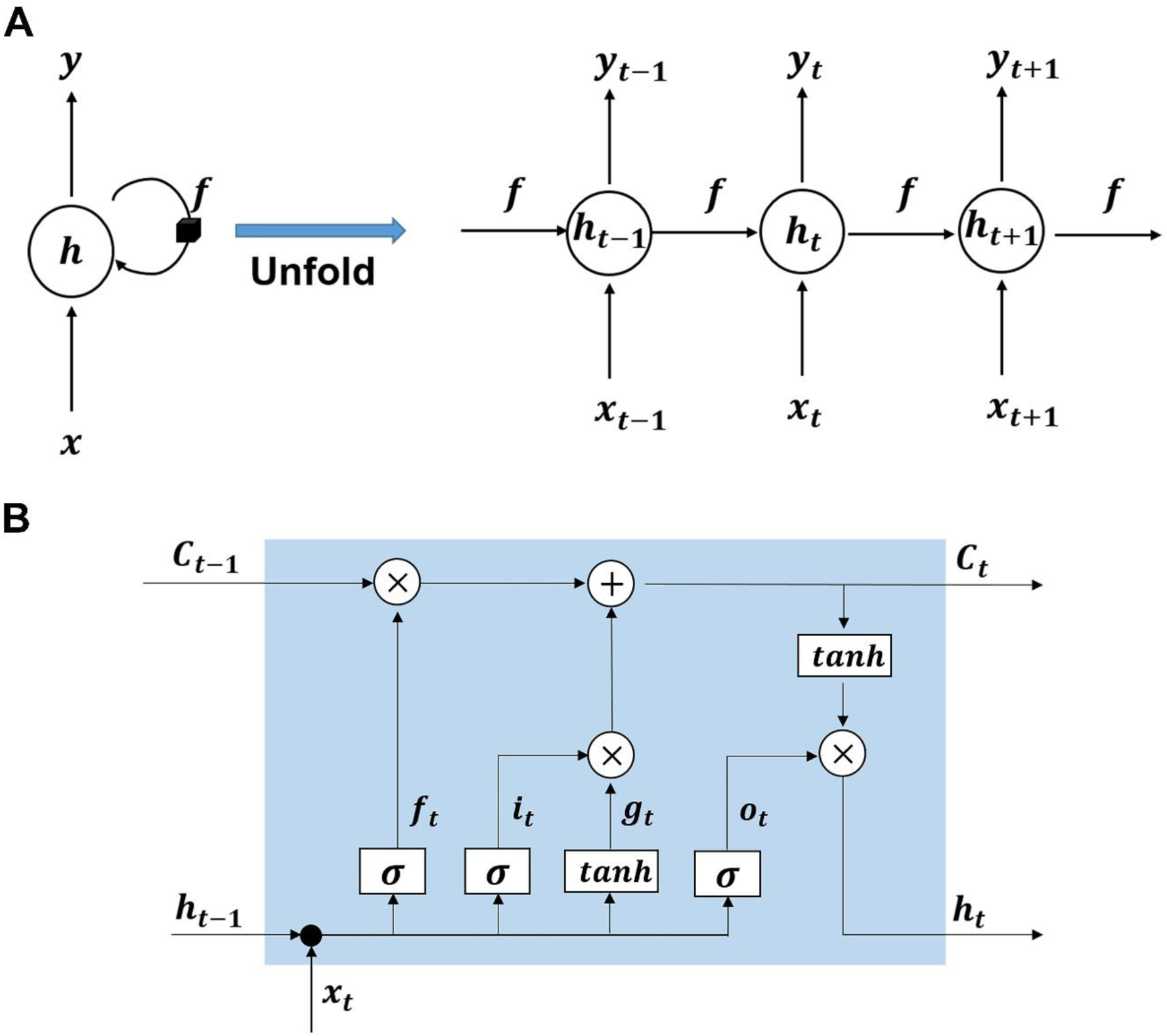
The architecture of RNN and LSTM. **a** The RNN architecture, unrolled, and *f* is a recursive activation function. **b** The corresponding diagram of an LSTM architecture.

The key to LSTMs is the cell state that enables the network to have a memory of past events, which determines the output at the current step and influences the next cell state. The dynamic variations of an LSTM architecture from the previous state *h*_*t*−1_ to current state *h*_*t*_ is performed by composite function below and the corresponding diagram was described in Figure 5B:

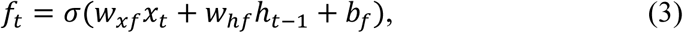

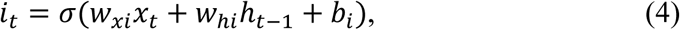

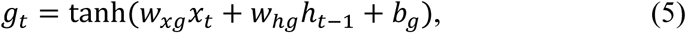

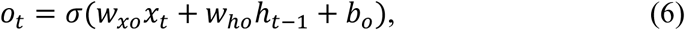

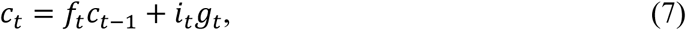

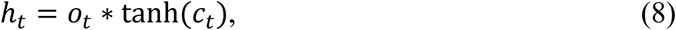

where *x*_*t*_ is the input vector of length *d* at the step *t, c*_*t*−1_ and *c*_*t*_ are the memory state vectors of length *h* at the step *t* − 1 and *t, σ* is the logistic sigmoid function, and *f*_*t*_, *i*_*t*_, *g*_*t*_ and *o*_*t*_ denote the forget gate, input gate, candidate memory state and output gate, respectively. The parameters such as *w*_*x*_, *w*_*h*_ and *b* are trainable, which correspond to weight matrixes of sizes *d* × *h, h* × *h* and bias vector parameters of sizes *h* × 1, respectively. More in-depth details about the LSTM architecture can be referred to the articles by Greff^53^ and Hochreiter et al^54^.

The model was trained with backpropagation through time (BPTT) by the ADAM optimizer, where the learning rate was set to 0.001 and the gradient norm clipping was 5 to avoid exploding gradients. The RNN model was implemented using TensorFlow (v1.2) and Keras (v2.0) in Python (v3.6), with the source code provided by Segler et al^30^.

### 4.3. DNN data set construction and processing

First, protein-ligand interaction data was downloaded from the GLASS database (http://zhanglab.ccmb.med.umich.edu/GLASS/), which contains 342539 ligands, 825 human GPCR proteins and 562871 unique GPCR-ligand entries. Subsequently, the IC_50_, EC_50_, and K_i_ data for the D_1_, D_2_, 5-HT_1A_ and 5-HT_2A_ receptors were selected and collected, along with their ligands. The SMILES strings and the target-associated bioactivities of these ligands were collected as the modeling data set. To expand the modeling data set, the experimental data sets, including K_i_, IC_50_ and EC_50_ data sets, for four targets were downloaded from Reaxys (a web-based tool for the retrieval of chemical information and data from published literature, journals and patents). Table 3 gives the detailed descriptions and the amounts of the four GPCR data sets used in the multitask DNN modeling.

We used the p-bioactivity, which is defined as − log_10_ *y*, throughout the modeling, where *y* is the raw bioactivity measured by K_i_, EC_50_, and IC_50_. Thus, the p-bioactivities ranged of from 0 to 14, where ligands with smaller values have higher activities. The ligands with multiple p-bioactivities were collected if there were considerable differences, data screening and cleaning were adopted, and the mean values were ultimately used.

### 4.4. MTDNN modeling and performance metrics

A typical neural network consists of multiple layers with neurons and acts as a transformation that maps an input feature vector to an output vector. Repeated linear and nonlinear transformations are performed on the inputs of these neurons, and backpropagation is used to update the weights of each layer. The transformation is given by the following:

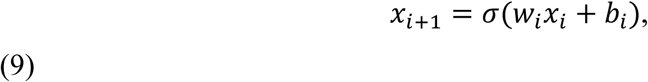

where *x*_*i*_ represents the input of the *i*-th layer; *w*_*i*_ and *b*_*i*_ represent the weight matrix and bias for the *i*-th layer, respectively; and *σ* is a nonlinearity activation functions using the rectified linear unit (ReLU). The learning process for updating the weights is conducted to reduce the difference between the predicted and observed activities. The difference given by the mean squared error (MSE) was used as the loss function for optimization. The fully connected feedforward neural network was used for backpropagation training, which has shown successful applications in multiple QSAR prediction tasks.^55^

The DNN architecture takes the molecular descriptors as inputs and produces a predicted value of p-bioactivity. The input molecular fingerprint based descriptor was ECFP4^56^, which was specifically designed for structure-activity modeling.^57^ Single-task DNNs have one output layer, while the MTDNN has multiple output layers. Here, the model has four output layers that produce the predicted bioactivities (pK_i_, pIC_50_ and pEC_50_) for the D_1_/D_2_/5-HT_1A_/5-HT_2A_ receptors. A random stratified classification was conducted to split the processed data into ten equal parts. One tenth of the data set was selected for evaluating the final model as an external test set. Eight tenths of the data were used as the training set, and the other tenth of the data set was used as the validation set to optimize the hyperparameters. In the training process, the optimizer, activation function and loss function were set as an ADAM optimizer, ReLU and MSE, respectively. To avoid overfitting, early stopping strategy was used where training was terminated when r^2^ on the validation set did not increase for 4 consecutive iterations.

r^2^ is defined as the square of the Pearson correlation coefficient between the predicted and observed activities in the test set, and this value ranges from 0 to 1 for all data sets,

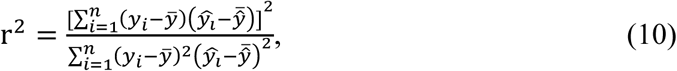

where *y*_*i*_ is the observed activity, 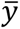 is the mean of the observed activities, *ŷ*_*i*_ is the predicted activity, 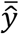 is the mean of the predicted activities, and n is the number of ligands. As the equation shows, better model performance results in values closer to 1.

The other metric used to evaluate prediction performance is MAE,

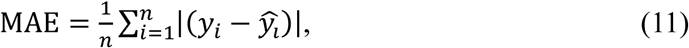

where *y*_*i*_ and *ŷ*_*i*_ are the observed and the predicted values, and n is the number of molecules in the data set. In this case, better model performance results in a smaller MAE value.

The DNN and MTDNN models were implemented using TensorFlow (v1.2) and Deepchem (v2.1) in Python (v3.6). All data are publicly available, with information about data location reported in the materials and methods.

### 4.5. Synthetic procedures

All commercially available materials and solvents were used without any further purification. TLC analyses were performed on Yantai Jiangyou silica gel HSGF254 plates. ^1^H NMR spectra and ^13^C NMR spectra were recorded at room temperature on a Bruker AVANCE-400 using TMS as an internal standard. Mass spectra were recorded on a Finnigan LTQ and 1290-6545 UHPLC-QTOF. The purity of test compounds were determined by HPLC and the purity of all test compounds is not less than 95%.

#### 4.5.1. Synthesis of 3

4-(4-(4-(benzo[b]thiophen-4-yl)piperazin-1-yl)butyl)-2H-benzo[b][1,4]oxazin- 3(4H)-one (**3**). Compound **3-a** (500 mg, 3.36 mmol) was dissolved in DMF (6 mL), NaH (60% in mineral oil, 270 mg, 6.72 mmol) was added in portions under ice bath followed by stirring for 1 h at 30 °C. Then, 1,4-dibromobutane (3.63 g, 16.8 mmol) was added followed by stirring at 30 °C overnight. The reaction mixture was extracted with ethyl acetate, washed three times with brine, dried, subjected to column chromatography using the mixture of petroleum etheracetone (40:1) as eluent to give **3-b** as a pale yellow oil (820 mg, yield 86%). Compound **3-b** (200 mg, 0.70 mmol), 1-(benzo[b]thiophen-4-yl)piperazine hydrochloride (195 mg, 0.77 mmol) and potassium carbonate (242 mg, 1.75 mmol) were added to acetonitrile (3 ml) under a nitrogen atmosphere and the mixture was stirred at reflux for 5 h. The reaction mixture was concentrated, washed three times with brine, dried, subjected to column chromatography to give **3** as a white solid (178 mg, 60%). 1H NMR (500 MHz, DMSO-*d*_6_) δ 7.69 (d, *J* = 5.5 Hz, 1H), 7.61 (d, *J* = 8.0 Hz, 1H), 7.39 (d, *J* = 5.5 Hz, 1H), 7.32 – 7.25 (m, 2H), 7.10 – 7.03 (m, 1H), 7.03 – 6.98 (m, 2H), 6.89 (d, *J* = 7.6 Hz, 1H), 4.63 (s, 2H), 3.94 (t, *J* = 7.4 Hz, 2H), 3.06 (s, 4H), 2.61 (s, 4H), 2.43 (s, 2H), 1.68 – 1.48 (m, 4H). ^13^C NMR (125 MHz, DMSO) δ 164.43, 146.95, 145.42, 141.10, 133.77, 128.61, 127.04, 125.53, 124.04, 123.26, 122.22, 118.21, 117.16, 115.89, 113.04, 67.48, 55.48, 51.55, 48.77, 24.49, 20.85. MS (ESI) m/z = 422.1 ([M+H]^+^). HRMS (ESI) m/z 422.1895 (calcd 422.1824 for C_24_H_28_N_3_O_2_S^+^ [M + H]^+^).

#### 4.5.2. Synthesis of 6

1-(4-(4-(benzo[b]thiophen-4-yl)piperazin-1-yl)butyl)-8-fluoro-3,4-dihydro-1H- benzo[e][1,4]diazepine-2,5-dione (**6**).The title compound was prepared in 30% yield by two steps from 8-fluoro-3,4-dihydro-1H-benzo[e][1,4]diazepine-2,5-dione and 1- (benzo[b]thiophen-4-yl)piperazine hydrochloride following the procedure described for synthesis of **3**. ^1^H NMR (300 MHz, DMSO-*d*_6_) δ 8.77 (t, *J* = 5.9 Hz, 1H), 7.75 (dd, *J* = 8.7, 6.7 Hz, 1H), 7.69 (d, *J* = 5.5 Hz, 1H), 7.61 (d, *J* = 8.0 Hz, 1H), 7.47 (dd, *J* = 11.0, 2.4 Hz, 1H), 7.37 (d, *J* = 5.5 Hz, 1H), 7.33 – 7.14 (m, 2H), 6.87 (d, *J* = 7.7 Hz, 1H), 4.27 (s, 1H), 3.85 – 3.59 (m, 2H), 3.57 – 3.40 (m, 1H), 3.00 (s, 4H), 2.37 – 2.17 (m, 2H), 1.60 – 1.14 (m, 4H). MS (ESI) *m/z* = 467.4 ([M+H]^+^). ^13^C NMR (125 MHz, DMSO) δ 169.75, 167.61, 164.28 (d, *J* = 248.7 Hz), 148.73, 141.81 (d, *J* = 10.8 Hz), 140.87, 133.85, 132.92 (d, *J* = 10.2 Hz), 126.56, 126.29, 125.58, 122.35, 117.10, 113.44 (d, *J* = 21.9 Hz), 112.48, 110.28 (d, *J* = 24.8 Hz), 57.32, 53.29, 52.13, 45.58, 45.41, 25.16, 23.23. HRMS (ESI) m/z 467.1923 (calcd 467.1839 for C_25_H_28_N_4_O_2_S^+^ [M + H]^+^).

#### 4.5.3. Synthesis of 7

1-(4-(4-(benzo[b]thiophen-4-yl)piperazin-1-yl)butyl)-7-hydroxy-3,4-dihydro-1H- benzo[c][1,2]thiazine 2,2-dioxide (**7**).The title compound was prepared in 21% yield by two steps from 7-hydroxy-3,4-dihydro-1H-benzo[c][1,2]thiazine 2,2-dioxide and 1- (benzo[b]thiophen-4-yl)piperazine hydrochloride following the procedure described for synthesis of **3**. ^1^H NMR (300 MHz, DMSO-*d*_6_) δ 9.48 (s, 1H), 7.69 (d, *J* = 5.5 Hz, 1H), 7.61 (d, *J* = 8.0 Hz, 1H), 7.39 (d, *J* = 5.5 Hz, 1H), 7.27 (t, *J* = 7.8 Hz, 1H), 7.00 (d, *J* = 8.2 Hz, 1H), 6.89 (d, *J* = 7.6 Hz, 1H), 6.52 (s, 1H), 6.47 (d, *J* = 8.4 Hz, 1H), 3.70 (t, *J* = 7.3 Hz, 2H), 3.40 (t, *J* = 7.0 Hz, 2H), 3.20 (t, *J* = 7.0 Hz, 2H), 3.07 (s, 4H), 2.61 (s, 4H), 2.41 (s, 2H), 1.77 – 1.46 (m, 4H). MS (ESI) *m/z* = 472.2 ([M+H]^+^). HRMS (ESI) m/z 472.1726 (calcd 472.1650 for C_24_H_30_N_3_O_3_S_2_^+^ [M + H]^+^).

#### 4.5.4. Synthesis of 8

1-(4-(4-(benzo[b]thiophen-4-yl)piperazin-1-yl)butyl)quinazoline-2,4(1H,3H)-dione (**8**).The title compound was prepared in 25% yield by two steps from quinazoline- 2,4(1H,3H)-dione and 1-(benzo[b]thiophen-4-yl)piperazine hydrochloride following the procedure described for synthesis of **3**. ^1^H NMR (300 MHz, DMSO-*d*_6_) δ 11.43 (s, 1H), 7.94 (d, *J* = 7.8 Hz, 1H), 7.75 – 7.56 (m, 3H), 7.43 – 7.35 (m, 1H), 7.34 – 7.13 (m, 3H), 6.88 (d, *J* = 7.7 Hz, 1H), 3.93 (t, *J* = 7.0 Hz, 2H), 3.05 (s, 4H), 2.59 (s, 4H), 2.40 (s, 2H), 1.77 – 1.41 (m, 4H). MS (ESI) *m/z* = 435.5 ([M+H]^+^). ^13^C NMR (125 MHz, DMSO) δ 162.39, 150.63, 148.71, 140.87, 139.87, 135.39, 133.85, 127.84, 126.28, 125.56, 122.93, 122.37, 117.10, 115.54, 114.25, 112.51, 57.97, 53.47, 52.13, 25.92, 24.27. HRMS (ESI) m/z 435.1839 (calcd 435.1776 for C_24_H_27_N_4_O_2_S^+^ [M + H]^+^).

#### 4.5.5. Synthesis of 9

1-(4-(4-(benzo[b]thiophen-4-yl)piperazin-1-yl)butyl)-7-methoxyquinoxalin-2(1H)- one (**9**).The title compound was prepared in 15% yield by two steps from 7- methoxyquinoxalin-2(1H)-one and 1-(benzo[b]thiophen-4-yl)piperazine hydrochloride following the procedure described for synthesis of **3**. ^1^H NMR (300 MHz, DMSO-*d*_6_) δ 8.05 (s, 1H), 7.76 (d, *J* = 9.0 Hz, 1H), 7.69 (d, *J* = 5.3 Hz, 1H), 7.61 (d, *J* = 8.1 Hz, 1H), 7.39 (d, *J* = 5.3 Hz, 1H), 7.28 (t, *J* = 7.7 Hz, 1H), 7.11 (s, 1H), 7.02 (d, *J* = 9.0 Hz, 1H), 6.89 (d, *J* = 7.4 Hz, 1H), 4.24 (t, *J* = 6.9 Hz, 2H), 3.90 (s, 3H), 3.07 (s, 4H), 2.61 (s, 4H), 2.44 (s, 2H), 1.80 – 1.53 (m, 4H). MS (ESI) *m/z* = 449.4 ([M+H]^+^). ^13^C NMR (125 MHz, DMSO) δ 161.91, 154.86, 148.71, 146.88, 140.89, 134.23, 133.86, 131.77, 128.40, 126.24, 125.53, 122.35, 117.08, 112.46, 111.00, 99.36, 57.32, 56.35, 53.45, 52.07, 41.46, 24.67, 23.71. HRMS (ESI) m/z 449.2005 (calcd 449.1933 for C_25_H_29_N_4_O_2_S^+^ [M + H]^+^).

#### 4.5.6. Synthesis of 10

7-amino-1-(4-(4-(benzo[b]thiophen-4-yl)piperazin-1-yl)butyl)-3,4-dihydro-1H- benzo[c][1,2]thiazine 2,2-dioxide (**10**).The title compound was prepared in 42% yield by two steps from 7-amino-3,4-dihydro-1H-benzo[c][1,2]thiazine 2,2-dioxide and 1- (benzo[b]thiophen-4-yl)piperazine hydrochloride following the procedure described for synthesis of **3**. ^1^H NMR (300 MHz, DMSO-*d*_6_) δ 7.69 (d, *J* = 5.5 Hz, 1H), 7.61 (d, *J* = 7.5 Hz, 1H), 7.39 (d, *J* = 5.5 Hz, 1H), 7.27 (t, *J* = 7.9 Hz, 1H), 6.90 (d, *J* = 7.9 Hz, 1H), 6.84 (d, *J* = 7.8 Hz, 1H), 6.36 – 6.25 (m, 2H), 5.10 (s, 2H), 3.64 (d, *J* = 9.1 Hz, 2H), 3.33 (s, 2H), 3.20 – 3.00 (m, 6H), 2.61 (s, 4H), 2.45 – 2.35 (m, 2H), 1.75 – 1.43 (m, 4H). ^13^C NMR (125 MHz, DMSO) δ 148.65, 141.00, 140.88, 133.86, 130.43, 126.32, 125.58, 122.37, 117.13, 112.55, 110.89, 110.04, 103.84, 57.76, 53.42, 52.15, 46.92, 46.01, 27.14, 27.02. MS (ESI) *m/z* = 471.5 ([M+H]^+^). HRMS (ESI) m/z 471.1878 (calcd 471.1810 for C_24_H_31_N_4_O_2_S_2_+ [M + H]^+^).

#### 4.5.7. Synthesis of 11

1-(4-(4-(benzo[b]thiophen-4-yl)piperazin-1-yl)butyl)-4-methyl-3,4-dihydro-1H- benzo[e][1,4]diazepine-2,5-dione hydrochloride (**11**). Compound **11-a** was prepared in 24% yield by two steps from 3,4-dihydro-1H-benzo[e][1,4]diazepine-2,5-dione and 1- (benzo[b]thiophen-4-yl)piperazine hydrochloride following the procedure described for synthesis of **3**. ^1^H NMR (300 MHz, Chloroform-*d*) δ 7.87 (dd, *J* = 7.7, 1.5 Hz, 1H), 7.62 – 7.53 (m, 2H), 7.42 – 7.21 (m, 5H), 7.08 (t, *J* = 7.3 Hz, 2H), 6.89 (d, *J* = 7.7 Hz, 1H), 4.44 – 4.25 (m, 1H), 3.79 (dd, *J* = 8.6, 6.2 Hz, 2H), 3.64 (d, *J* = 13.4 Hz, 1H), 3.27 (s, 4H), 2.84 (s, 4H), 2.57 (s, 2H), 1.59 (dd, *J* = 12.9, 6.4 Hz, 4H). MS (ESI) *m/z* = 449.3 ([M+H]^+^). **11-a** (90 mg, 0.20 mmol) was dissolved in DMF (3 mL), NaH (60% in mineral oil, 9 mg, 0.22 mmol) was added under ice bath followed by stirring under ice bath for 30 min. Dimethyl sulfate (21 mL, 0.22 mmol) was added under ice bath followed by stirring at room temperature overnight. The reaction mixture was poured into ice water and extracted with EA. The combined organic phase was washed twice with brine, dried over anhydrous sodium sulfate, subjected to column chromatography using DCM:MeOH (100:1–20:1) as eluent to give a crude product. The crude residue was dissolved in ethanol, hydrogen chloride-ethanol solution was added thereto under stirring, the resulting salt was slurried in MTBE/MeOH system, filtered and dried to give **11** as a light yellow solid (14 mg, yield 14%). ^1^H NMR (300 MHz, DMSO-*d*_6_) δ 10.69 (s, 1H), 7.77 (d, *J* = 5.5 Hz, 1H), 7.70 (d, *J* = 7.8 Hz, 2H), 7.67 – 7.45 (m, 3H), 7.41 – 7.27 (m, 2H), 6.96 (d, *J* = 7.6 Hz, 1H), 4.31 – 4.12 (m, 1H), 4.05 (d, *J* = 14.8 Hz, 1H), 3.83 – 3.63 (m, 2H), 3.61 – 3.42 (m, 4H), 3.30 – 2.99 (m, 9H), 1.72 – 1.32 (m, 4H). ^13^C NMR (125 MHz, DMSO) δ 168.39, 166.91, 146.98, 141.09, 139.55, 133.81, 132.49, 130.45, 130.25, 127.05, 126.19, 125.53, 122.79, 122.17, 118.22, 113.05, 52.97, 51.70, 48.89, 45.52, 35.66, 25.03. MS (ESI) *m/z* = 463.4 ([M+H]^+^). HRMS (ESI) m/z 463.2162 (calcd 462.2089 for C_26_H_31_N_4_O_2_S^+^ [M + H]^+^).

### 4.6. Functional Activity Assays

All the compounds were screened in 5-HT_1A_ agonist and D_2_ antagonist mode assays using Ultra Lance and 5-HT_2A_ antagonist mode assays using FLIPR.

IC_50_/EC_50_ values were calculated from the concentration–inhibition curves by nonlinear regression analysis using GraphPad Prism 6.0. Ultra Lance cAMP assay: (1) Transfer compound to assay plate by Echo; (2) Collect cells with stimulation buffer; (3) Reaction: 1) transfer 10 μL of cell solution to assay plate, 2) centrifuge at 600 rpm for 3 min and incubate for 60 min at room temperature, 3) add 5 μL of 4X Eu-cAMP trace solution and 5 μL of 4X ULight^™^-anti-cAMP solution to assay plate, 4) centrifuge at 600 rpm for 3 min and incubate for 60 min at room temperature; (4) Read using an Envision plate reader.

FLIPR assays: (1) Seed cells at a density of 10 K cell/well, and incubate the cells at 37 °C under 5% CO _2_ for 16–24 h; (2) Load cells with 30 μL of calcium 5, and incubate the samples at 37 °C under 5% CO_2_ for 1 h; (3) Transfer compounds to a compound plate with 30 μL of assay buffer by Echo; (4) Add 15 μL of the test compound to each well and incubate for 10 min at room temperature; (5) Add 22.5 μL of inducer to each well and measure the calcium flux signal with FLIPR.

### 4.7. PCP-induced Hyperlocomotion

All procedures performed on animals were in accordance with regulations and established guidelines, and were reviewed and approved by the Institutional Animal Care and Use Committee at Shanghai Institute of Materia Medica, Chinese Academy of Sciences (Shanghai, China).

Male ICR mice (28∼33 g, 8 ∼ 10 mice in each group) were used. Animals were individually placed into the locomotor activity chamber, and spontaneous activities were measured for 30 min after intragastric administration of the test compounds (3 and 10 mg/kg) or aripiprazole (3 mg/kg). Then, the animals were administered PCP (7 mg/kg, 10 mL/kg, ip), and placed back into the experimental chamber. The animals were habituated for 10 min before the 60 min measurement period. The results are expressed as the mean ± SEM of the distance traveled. Statistical evaluations were performed by one-way ANOVA followed by Dunnett’s post hoc test.

Catalepsy Test. Mice were orally dosed with vehicle or compounds. Catalepsy was evaluated on a metal bar 0.3 cm in diameter positioned 4.5 cm above the tabletop. The test consisted in positioning the animal with its forepaws on the bar and recording how long it remained hanging onto the bar. A mean immobility score of 20 s was used as the criterion for the presence of catalepsy.

## Supporting information

Supplemental materials, and will be used for the link to the file on the preprint site.

## Acknowledgements

We thank Prof. Mark P. Waller at Shanghai University for kindly providing codes of molecular generation and helpful discussions. This project is supported by the National Natural Science Foundation of China (81773634 to M.Z. and 81703338 to Y.H.), National Science & Technology Major Project “Key New Drug Creation and Manufacturing Program”, China (Number: 2018ZX09711002 to H.J.), and “Personalized Medicines—Molecular Signature-based Drug Discovery and Development”, Strategic Priority Research Pro-gram of the Chinese Academy of Sciences (XDA12050201 to M.Z. and XDA12040331 to Y.H.).

## Author Contributions

H.J., M.Z. and J.S. conceived the project. X.T. wrote code and conducted computational analysis of deep learning models. X.J., Y.H. and Z.W. synthesized the compounds and carried out the in vitro and in vivo experiments. X.T., X.J., Y.H. wrote the paper. F.Z., X.L., Z.X., Z.L., C.C. and X.L. collected and analysed the data. Q.Z., Y.X., F.Y. and C.W. performed the chemical synthesis. All authors discussed the results and commented on the manuscript.

## Appendix A

Supplementary data Supplementary data to this article can be found online.

## References

[1] H.M. Ibrahim; C.A. Tamminga Schizophrenia: Treatment Targets Beyond Monoamine Systems. Annu. Rev. Pharmacol. Toxicol. 51 (2011) 189–209.

[2] B.L. Roth; D.J. Sheffler; W.K. Kroeze Magic shotguns versus magic bullets: selectively non-selective drugs for mood disorders and schizophrenia. Nat. Rev. Drug Discovery 3 (2004) 353.

[3] R. Tandon; H.A. Nasrallah; M.S. Keshavan Schizophrenia, “just the facts” 4. Clinical features and conceptualization. Schizophr. Res. 110 (2009) 1–23.

[4] S.R. Marder; W.C. Wirshing; T. Van Putten Drug treatment of schizophrenia: overview of recent research. Schizophr. Res. 4 (1991) 81–90.

[5] S. Miyamoto; N. Miyake; L. Jarskog; W. Fleischhacker; J. Lieberman Pharmacological treatment of schizophrenia: a critical review of the pharmacology and clinical effects of current and future therapeutic agents. Mol Psychiatry 17 (2012) 1206.

[6] P.M. Haddad; S.G. Sharma Adverse effects of atypical antipsychotics. CNS Drugs 21 (2007) 911–936.

[7] M. De Hert; J. Detraux; R. van Winkel; W. Yu; C.U. Correll Metabolic and cardiovascular adverse effects associated with antipsychotic drugs. Nat. Rev. Endocrinol. 8 (2011) 114.

[8] G. Gründer Cariprazine, an orally active D2/D3 receptor antagonist, for the potential treatment of schizophrenia, bipolar mania and depression. Curr. Opin. Investig. Drugs. 11 (2010) 823–832.

[9] S.H. Schultz; S.W. North; C.G. Shields Schizophrenia: a review. Am. Fam. Phys. 75 (2007) 1821–9.

[10] B.J. Kinon; J.A. Lieberman Mechanisms of action of atypical antipsychotic drugs: a critical analysis. Psychopharmacology 124 (1996) 2–34.

[11] H.-J. Möller Management of the Negative Symptoms of Schizophrenia. CNS Drugs 17 (2003) 793–823.

[12] R. Schreiber; A. Newman-Tancredi Improving cognition in schizophrenia with antipsychotics that elicit neurogenesis through 5-HT(1A) receptor activation. Neurobiol Learn Mem 110 (2014) 72–80.

[13] P. Celada; A. Bortolozzi; F. Artigas Serotonin 5-HT1A Receptors as Targets for Agents to Treat Psychiatric Disorders: Rationale and Current Status of Research. CNS Drugs 27 (2013) 703–716.

[14] D.A. Shapiro; S. Renock; E. Arrington; L.A. Chiodo; L.X. Liu; D.R. Sibley; B.L. Roth; R. Mailman Aripiprazole, a novel atypical antipsychotic drug with a unique and robust pharmacology. Neuropsychopharmacology 28 (2003) 1400–11.

[15] K. Maeda; H. Sugino; H. Akazawa; N. Amada; J. Shimada; T. Futamura; H. Yamashita; N. Ito; R.D. McQuade; A. Mørk Brexpiprazole I: in vitro and in vivo characterization of a novel serotonin-dopamine activity modulator. J. Pharmacol. Exp. Ther. 350 (2014) 589–604.

[16] B. Kiss; A. Horváth; Z. NÉmethy; É. Schmidt; I. Laszlovszky; G. Bugovics; K. Fazekas; K. Hornok; S. Orosz; I. Gyertyán Cariprazine (RGH-188), a dopamine D3 receptor-preferring, D3/D2 dopamine receptor antagonist–partial agonist antipsychotic candidate: in vitro and neurochemical profile. J. Pharmacol. Exp. Ther. 333 (2010) 328–340.

[17] P.G. Polishchuk; T.I. Madzhidov; A. Varnek Estimation of the size of drug-like chemical space based on GDB-17 data. J. Comput.-Aided Mol. Des. 27 (2013) 675–679.

[18] S. Paricharak; O. MÉndez-Lucio; A. Chavan Ravindranath; A. Bender; A.P. Ijzerman; G.J.P. van Westen Data-driven approaches used for compound library design, hit triage and bioactivity modeling in high-throughput screening. Brief Bioinform. (2016) bbw105.

[19] O.M. Becker; Y. Marantz; S. Shacham; B. Inbal; A. Heifetz; O. Kalid; S. Bar-Haim; D. Warshaviak; M. Fichman; S. Noiman G protein-coupled receptors: in silico drug discovery in 3D. Proc. Natl. Acad. Sci. U. S. A. 101 (2004) 11304–11309.

[20] D. Wootten; A. Christopoulos; P.M. Sexton Emerging paradigms in GPCR allostery: implications for drug discovery. Nat. Rev. Drug Discovery 12 (2013) 630.

[21] F. Dey; A. Caflisch Fragment-based de novo ligand design by multiobjective evolutionary optimization. J. Chem. Inf. Model. 48 (2008) 679–690.

[22] C.A. Nicolaou; N. Brown; C.S. Pattichis Molecular optimization using computational multi-objective methods. Curr. Opin. Drug Discovery Dev. 10 (2007) 316–324.

[23] M. Olivecrona; T. Blaschke; O. Engkvist; H. Chen Molecular de-novo design through deep reinforcement learning. J. Cheminf. 9 (2017) 48.

[24] M. Awale; F. Sirockin; N. Stiefl; J.L. Reymond Drug analogs from fragment-based long short-term memory generative neural networks. J. Chem. Inf. Model. 59 (2019) 1347–1356.

[25] H. Altae-Tran; B. Ramsundar; A.S. Pappu; V. Pande Low data drug discovery with one-shot learning. ACS Cent. Sci. 3 (2017) 283–293.

[26] R. Gómez-Bombarelli; J.N. Wei; D. Duvenaud; J.M. Hernández-Lobato; B. Sánchez-Lengeling; D. Sheberla; J. Aguilera-Iparraguirre; T.D. Hirzel; R.P. Adams; A. Aspuru-Guzik Automatic chemical design using a data-driven continuous representation of molecules. ACS Cent. Sci. 4 (2018) 268–276.

[27] M. Popova; O. Isayev; A. Tropsha Deep reinforcement learning for de novo drug design. Sci. Adv. 4 (2018) eaap7885.

[28] A. Zhavoronkov; Y.A. Ivanenkov; A. Aliper; M.S. Veselov; V.A. Aladinskiy; A.V. Aladinskaya; V.A. Terentiev; D.A. Polykovskiy; M.D. Kuznetsov; A. Asadulaev Deep learning enables rapid identification of potent DDR1 kinase inhibitors. Nat Biotechnol (2019) 1–4.

[29] Y. Xu; J. Ma; A. Liaw; R.P. Sheridan; V. Svetnik Demystifying multitask deep neural networks for quantitative structure-activity relationships. J. Chem. Inf. Model. 57 (2017) 2490–2504.

[30] M.H. Segler; T. Kogej; C. Tyrchan; M.P. Waller Generating focused molecule libraries for drug discovery with recurrent neural networks. ACS Cent. Sci. 4 (2018) 120–131.

[31] P. Ertl; B. Rohde; P. Selzer Fast calculation of molecular polar surface area as a sum of fragment-based contributions and its application to the prediction of drug transport properties. J. Med. Chem. 43 (2000) 3714–3717.

[32] L.v.d. Maaten; G. Hinton Visualizing data using t-SNE. J. Mach. Learn. Res. 9 (2008) 2579–2605.

[33] S. Jafari; F. Fernandez-Enright; X.F. Huang Structural contributions of antipsychotic drugs to their therapeutic profiles and metabolic side effects. J. Neurochem. 120 (2012) 371–384.

[34] P. Ertl; A. Schuffenhauer Estimation of synthetic accessibility score of drug-like molecules based on molecular complexity and fragment contributions. J. Cheminf. 1 (2009) 8.

[35] R. Kadam; N. Roy Recent trends in drug-likeness prediction: a comprehensive review of in silico methods. Indian. J. Pharm. Sci. 69 (2007) 609–615.

[36] S. Tian; J. Wang; Y. Li; D. Li; L. Xu; T. Hou The application of in silico drug-likeness predictions in pharmaceutical research. Adv. Drug Delivery Rev. 86 (2015) 2–10.

[37] F. Chevillard; P. Kolb SCUBIDOO: a large yet screenable and easily searchable database of computationally created chemical compounds optimized toward high likelihood of synthetic tractability. J. Chem. Inf. Model. 55 (2015) 1824–1835.

[38] D. Merk; L. Friedrich; F. Grisoni; G. Schneider De novo design of bioactive small molecules by artificial intelligence. Mol. Inf. 37 (2018) 1700153.

[39] D. Merk; F. Grisoni; L. Friedrich; G. Schneider Tuning artificial intelligence on the de novo design of natural-product-inspired retinoid X receptor modulators. Nat. Commun. Chem. 1 (2018).

[40] Y. Li; L. Zhang; Z. Liu Multi-objective de novo drug design with conditional graph generative model. . Cheminf. 10 (2018) 33.

[41] H.Y. Meltzer What’s atypical about atypical antipsychotic drugs? Curr. Opin. Pharmacol. 4 (2004) 53–7.

[42] A. Newman-Tancredi; M.S. Kleven Comparative pharmacology of antipsychotics possessing combined dopamine D2 and serotonin 5-HT1A receptor properties. Psychopharmacology 216 (2011) 451–473.

[43] J.B. Baell; G.A. Holloway New substructure filters for removal of pan assay interference compounds (PAINS) from screening libraries and for their exclusion in bioassays. J. Med. Chem. 53 (2010) 2719–2740.

[44] A.K. Ghose; T. Herbertz; R.L. Hudkins; B.D. Dorsey; J.P. Mallamo Knowledge-based, central nervous system (CNS) lead selection and lead optimization for CNS drug discovery. ACS Chem. Neurosci. 3 (2011) 50–68.

[45] A. Reichel Addressing central nervous system (CNS) penetration in drug discovery: basics and implications of the evolving new concept. Chem. Biodivers. 6 (2009) 2030–2049.

[46] D.S. Johnson; C. Choi; L.K. Fay; D.A. Favor; J.T. Repine; A.D. White; H.C. Akunne; L. Fitzgerald; K. Nicholls; B.J. Snyder Discovery of PF-00217830: Arylpiperazine napthyridinones as D2 partial agonists for schizophrenia and bipolar disorder. Bioorg. Med. Chem. 21 (2011) 2621–2625.

[47] Y. Chen; S. Wang; X. Xu; X. Liu; M. Yu; S. Zhao; S. Liu; Y. Qiu; T. Zhang; B.-F. Liu Synthesis and biological investigation of coumarin piperazine (piperidine) derivatives as potential multireceptor atypical antipsychotics. J. Med. Chem. 56 (2013) 4671–4690.

[48] A. Czopek; M. Kołaczkowski; A. Bucki; H. Byrtus; M. Pawłowski; G. Kazek; A.J. Bojarski; A. Piaskowska; J. Kalinowska-Tłuścik; A. Partyka Novel spirohydantoin derivative as a potent multireceptor-active antipsychotic and antidepressant agent. Bioorg. Med. Chem. 23 (2015) 3436–3447.

[49] E.D. French; C. Pilapil; R. Quirion Phencyclidine binding sites in the nucleus accumbens and phencyclidine-induced hyperactivity are decreased following lesions of the mesolimbic dopamine system. Eur. J. Pharmacol. 116 (1985) 1–9.

[50] D.C. Javitt; S.R. Zukin Recent advances in the phencyclidine model of schizophrenia. Am. J. Psychiatry. 148 (1991) 1301.

[51] B.J. Morris; S.M. Cochran; J.A. Pratt PCP: from pharmacology to modelling schizophrenia. Curr. Opin. Pharmacol. 5 (2005) 101–106.

[52] J.L. Moreno; J. González-Maeso Preclinical models of antipsychotic drug action. Int. J. Neuropsychopharmacol. 16 (2013) 2131–2144.

[53] K. Greff; R.K. Srivastava; J. Koutník; B.R. Steunebrink; J. Schmidhuber LSTM: a search space odyssey. IEEE transactions on neural networks and learning systems 28 (2016) 2222–2232.

[54] S. Hochreiter; J. Schmidhuber Long short-term memory. Neural Comput 9 (1997) 1735–1780.

[55] J. Ma; R.P. Sheridan; A. Liaw; G.E. Dahl; V. Svetnik Deep neural nets as a method for quantitative structure–activity relationships. J. Chem. Inf. Model. 55 (2015) 263–274.

[56] H. Morgan The generation of a unique machine description for chemical structures-a technique developed at chemical abstracts service. J. Chem. Doc. 5 (1965) 107–113.

[57] D. Rogers; M. Hahn Extended-connectivity fingerprints. J. Chem. Inf. Model. 50 (2010) 742–754.

