## Supplemental materials, and will be used for the link to the file on the preprint site. for "Automated design and optimization of multitarget schizophrenia drug candidates by deep learning"

Contents of Supporting Information:

Figure S1. The homology correlations between dopaminergic and serotoninergic receptors.
Figure S2. The loss curves of pretrained and fine-tuned model
Figure S3. Random examples of the generated molecules from pretrained RNN model.
Figure S4. QED scores distributions and random samples of generated molecules, distribution of properties for comparing the generated molecules and the molecules exsiting in training set.
Figure S5. Distribution of properties for molecules designed by retrained language model and training set: histogram of training set (purple) and generated molecules (green) for (A) MolWt, (B) QED, (C) LogP, (D) SA.
Table S1. Molecules that predicted not active to GPCRs target.
Figure S6. Effects of compound 6, 7, 8, 9, 10, 11 and aripiprazole administered po on PCP-induced hyperactivity in mice; effects of compound 7, 10 and aripiprazole administered po on spontaneous activity in mice.

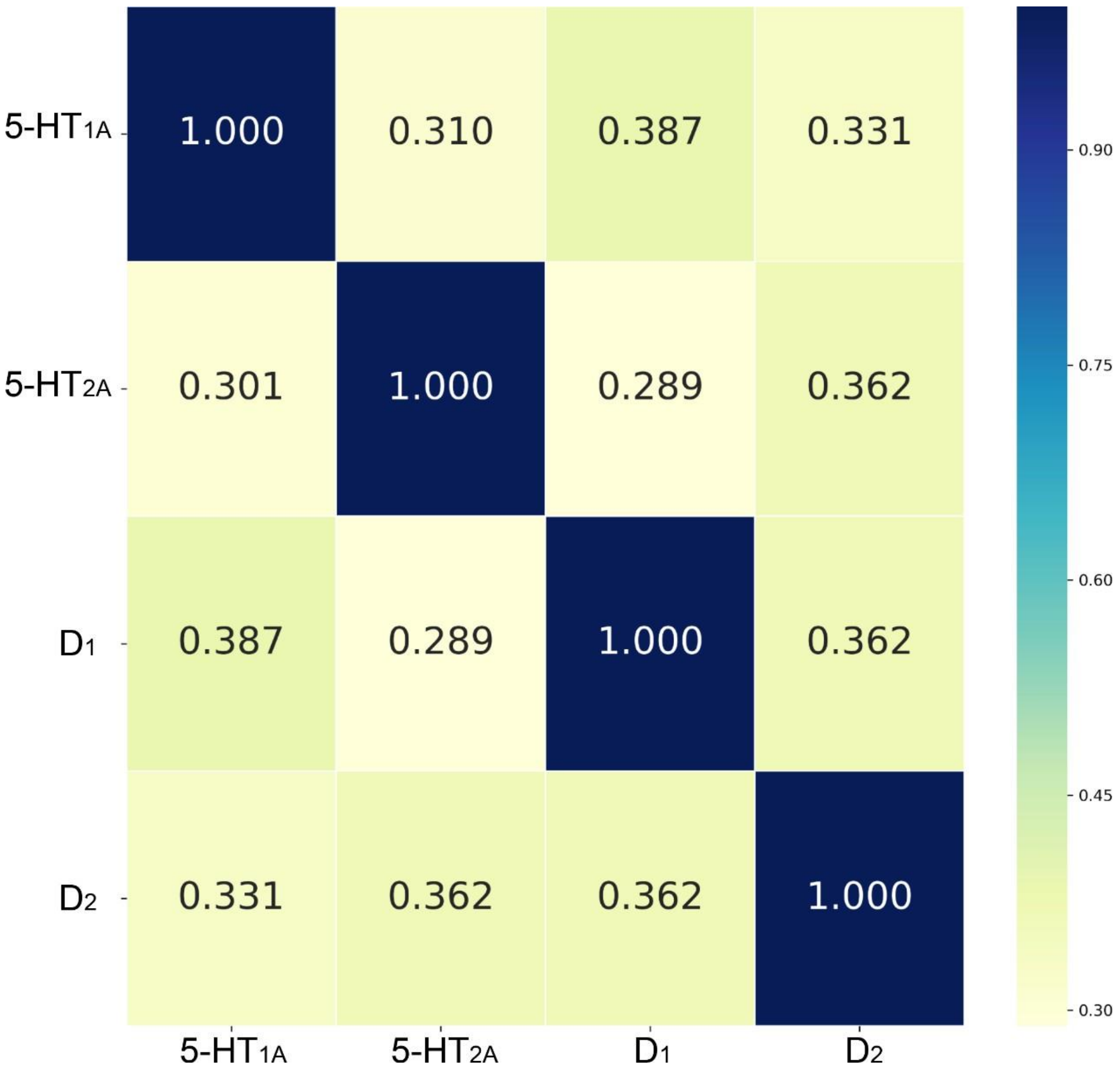


**Figure S1.** The homology correlation between dopaminergic and serotoninergic receptors.

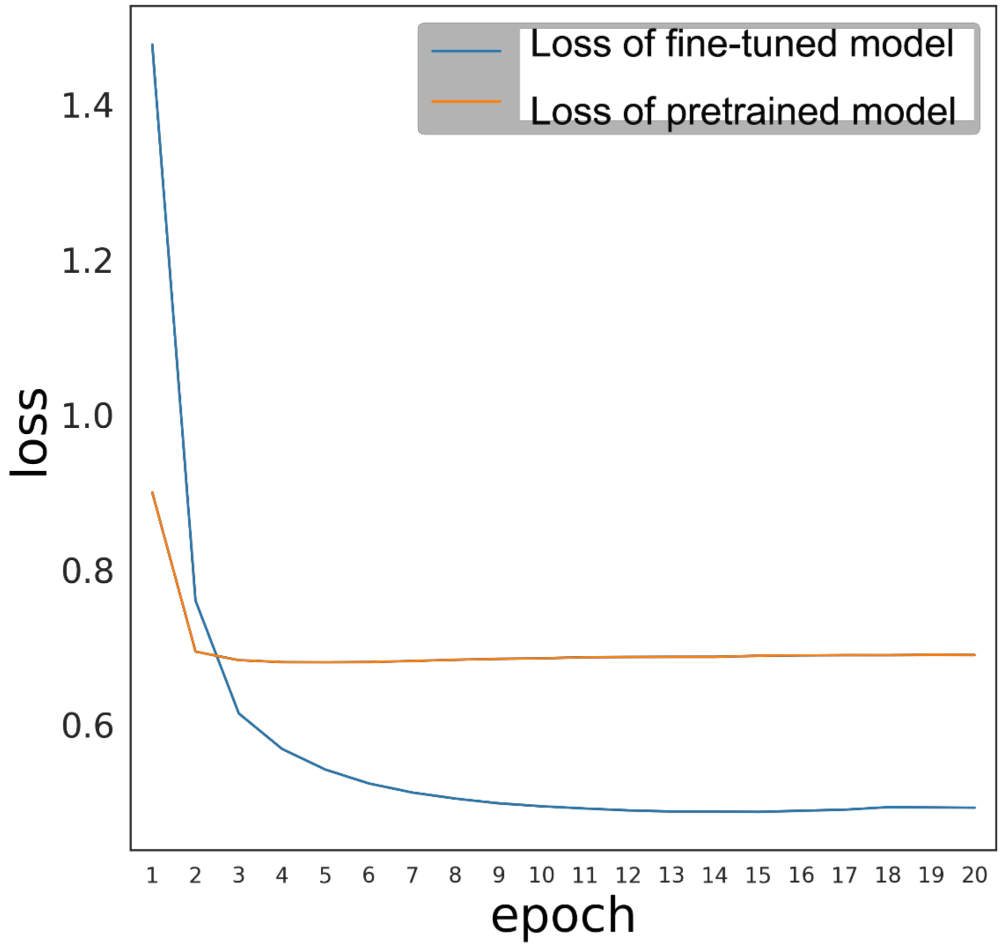


**Figure S2.** The loss curves of pretrained RNN model and fine-tuned RNN model.

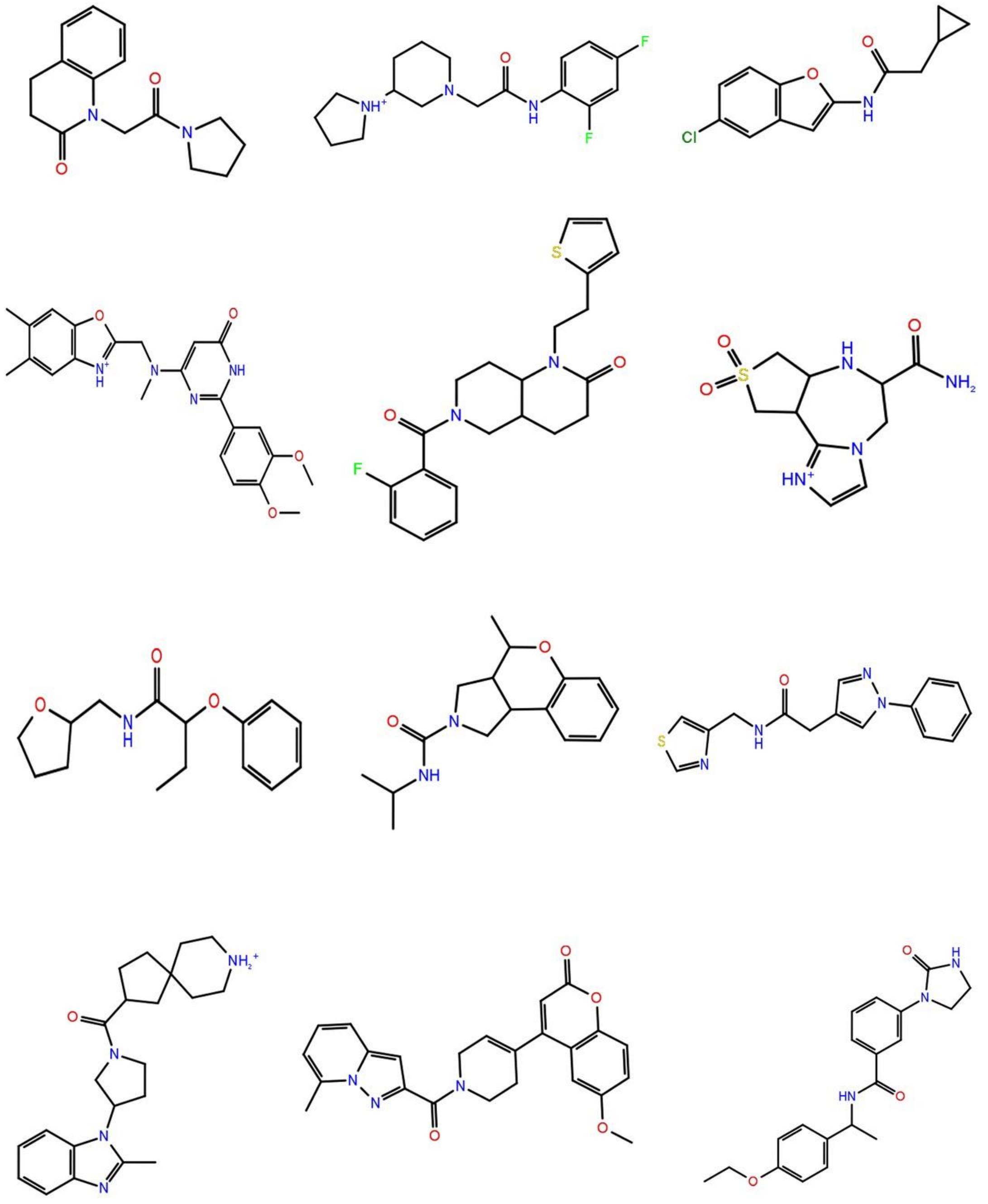

**Figure S3.** Random samples of generated molecules from the pretrained RNN model.

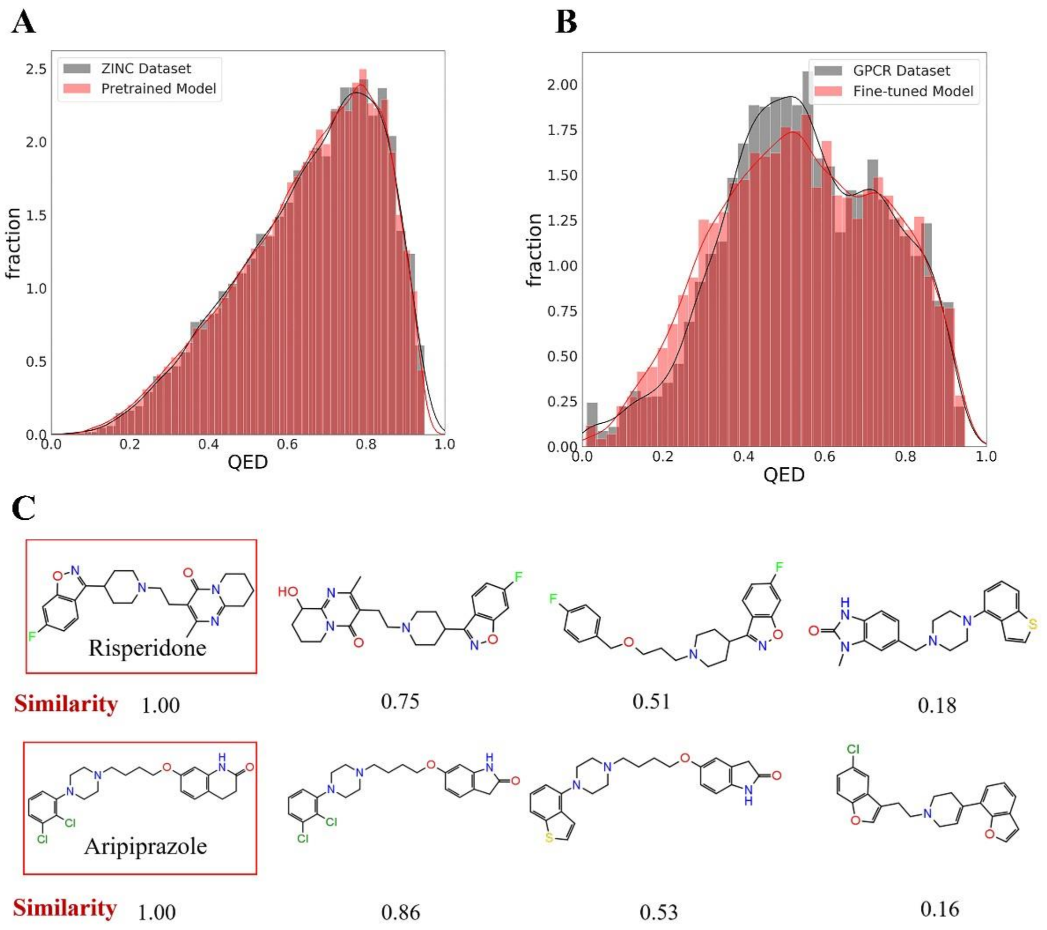


**Figure S4.** Distribution of the physicochemical properties for molecules generated with the fine-tuned model and random samples of the focused library. (A) Comparison of the QED scores of the molecules generated by the model pretrained with the *ZINC* data set. (B) Comparison of the QED scores of the molecules generated by the fine-tuned model trained with the published GPCR data set. (C) Samples of the molecules generated by the fine-tuned generative model.

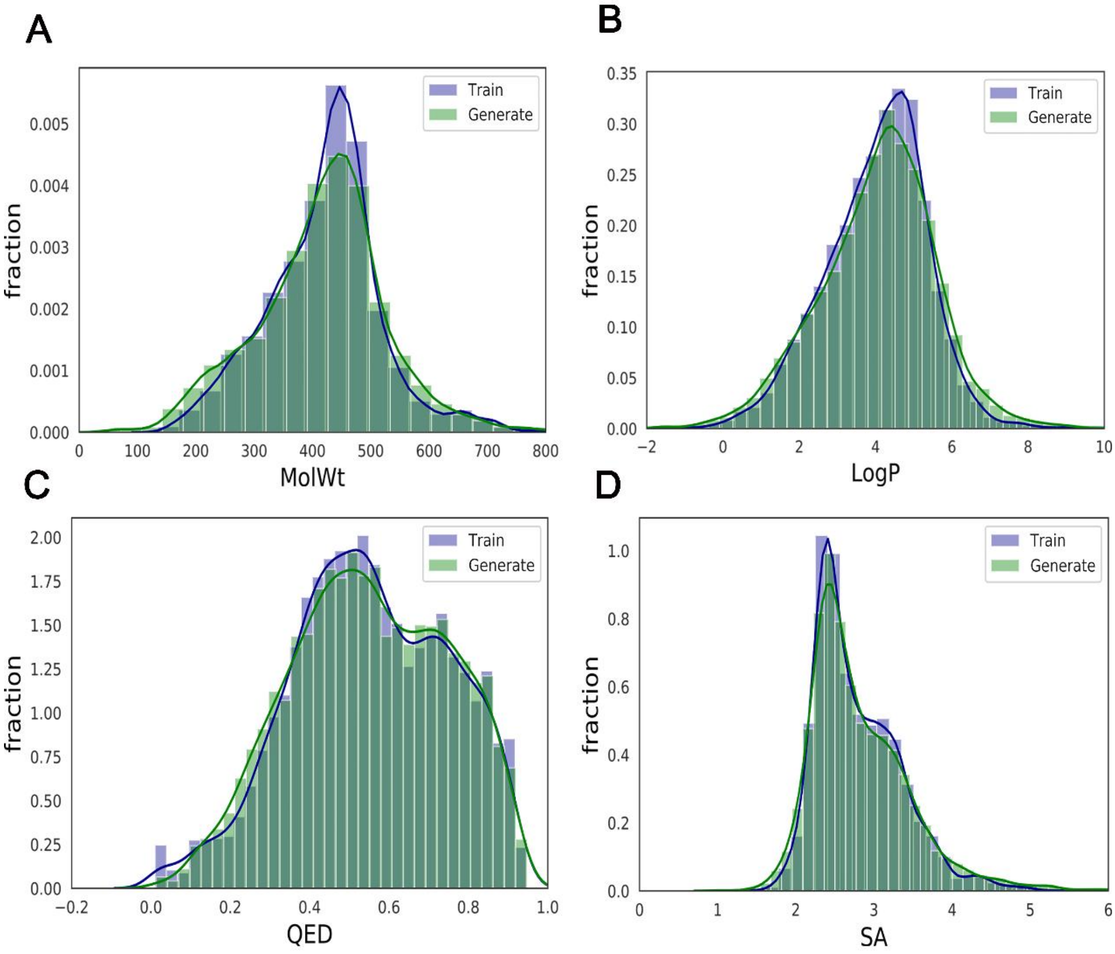


**Figure S5.** Distribution of properties (A) MolWt, (B) LogP, (C) QED, (D)SA for comparing the molecules designed by fine-tuned language model (green) and the molecules exsiting in training set (purple).

**Supplementary Table**

Table S1: Generated structures with imbalanced or less potent activities to GPCR targets.

| Compounds | Structure | Predicted 5-$HT_{1A}$ $pEC_{50}$ | Predicted $D_2$ $pIC_{50}$ | Predicted 5-$HT_{2A}$ $pIC_{50}$ |
|---|---|---|---|---|
| **12** | 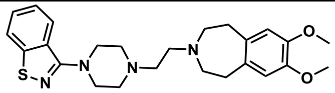 | 5.76 | 7.97 | 8.74 |
| **13** | 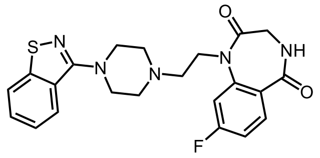 | 7.76 | 6.02 | 6.51 |
| **14** | 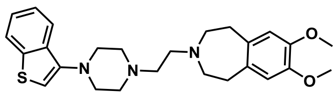 | 4.61 | 7.19 | 7.92 |
| **15** | 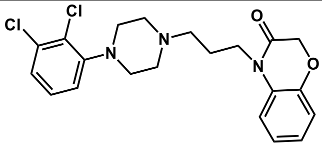 | 7.03 | 7.09 | 6.22 |

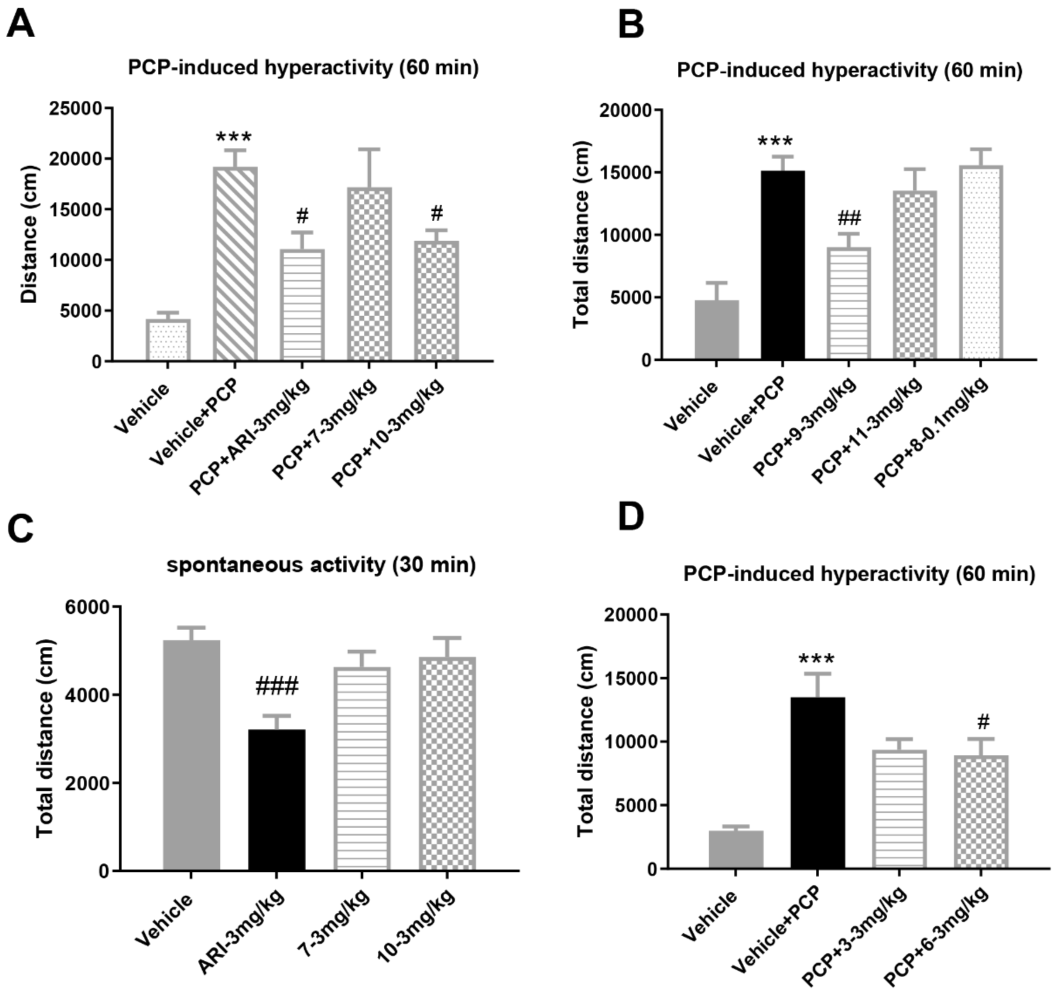


**Figure S6.** (A), (B), (C) Effects of compound **6, 7, 8, 9, 10** and **11** administered po on PCP-induced hyperactivity in mice (PCP: 7 mg/kg, ip). Locomotor activities were measured for 60 min following PCP administration, and the results are expressed as the mean ± SEM of the distance traveled. (D) Effects of aripiprazole (ARI, 3 mg/kg), 7 and 10 administered po on spontaneous activity in mice. The locomotor activities were measured during the 30 min following drug administration, and the results are expressed as the mean ± SEM of distance traveled. Statistical evaluation was performed by one-way ANOVA followed by Dunnett's post hoc test. #$p < 0.05$ versus PCP treatment; ##$p < 0.01$ versus PCP treatment; ***$p < 0.001$ versus vehicle treatment. ARI: aripiprazole.
